# Steady-State Regulation of COPII-Dependent Secretory Cargo Sorting by Inositol Trisphosphate Receptors, Calcium, and Penta EF Hand Proteins

**DOI:** 10.1101/2020.06.13.150144

**Authors:** Aaron Held, Jacob Lapka, John Sargeant, Jennet Hojanazarova, Corina Madreiter-Sokolowski, Roland Malli, Wolfgang F. Graier, Jesse C. Hay

## Abstract

Recently, we demonstrated that agonist-stimulated Ca^2+^ signaling involving IP3 receptors modulates ER export rates through activation of the penta-EF Hand (PEF) proteins apoptosis-linked gene-2 (ALG-2) and peflin. It is unknown, however, whether IP3Rs and PEF proteins regulate ER export rates at steady state. Here we tested this idea in normal rat kidney (NRK) epithelial cells by manipulation of IP3R isoform expression. Under standard growth conditions, spontaneous cytosolic Ca^2+^ oscillations occurred simultaneously in successive groups of contiguous cells, generating intercellular Ca^2+^ waves (ICWs) that moved across the monolayer periodically. Depletion of IP3R-3, typically the least promiscuous IP3R isoform, caused increased cell participation in ICWs in unstimulated cells. The increased spontaneous signaling was sufficient to cause increased ALG-2 and Sec31A, and decreased peflin localization at ER exit sites (ERES), resulting in increased ER-to-Golgi transport of the COPII client cargo VSV-G. The elevated ER-to-Golgi transport caused greater concentration of VSV-G at ERES and had reciprocal effects on transport of VSV-G and a bulk-flow cargo, though both cargos equally required Sec31A. Inactivation of client cargo sorting using 4-phenylbutyrate (4-PBA) had opposing reciprocal effects on client and bulk-flow cargo and neutralized any effect of ALG-2 activation on transport. This work extends our knowledge of ALG-2 mechanisms and indicates that in NRK cells, IP3R isoforms regulate homeostatic Ca^2+^ signaling that helps determine the basal secretion rate and stringency of COPII-dependent cargo sorting.

## INTRODUCTION

Calcium is a vital intracellular signaling molecule integral for a diverse array of physiological processes. Regulatory roles for Ca^2+^ in intracellular trafficking steps are still being elucidated. Recent work on ER-to-Golgi transport demonstrates involvement of luminal Ca^2+^ stores at a stage following cargo biogenesis and folding/assembly, apparently through the release of Ca^2+^ into the cytoplasm where it binds and activates the vesicle budding, docking and/or fusion machinery (1, 2). Effector mechanisms by which cytosolic Ca^2+^ modulates ER-to-Golgi transport appear to include penta-EF-hand-containing (PEF) protein adaptors that have been implicated in many Ca^2+^-dependent cellular phenomena (3). The PEF protein apoptosis-linked gene-2 (ALG-2) acts as a Ca^2+^ sensor at ER exit sites (ERES) and can stabilize the association of the COPII coat subunit Sec31A with the membrane when Ca^2+^ is present (4–7). However, regulation of ER cargo export by ALG-2 and Ca^2+^ does not follow the simple pattern suggested by these early studies. For one thing, sustained agonist-driven Ca^2+^ signaling results in a sharp ALG-2-dependent reduction of COPII targeting and ER export (8). For another, ALG-2 in cell extracts often exists in a stable heterodimer with another PEF protein–peflin–that binds ALG-2 in a Ca^2+^-inhibited manner (9, 10). The peflin-complexed form of ALG-2 appears to bind ERES, destabilize the COPII coat and suppress ER export, but through a distinct mechanism than during sustained Ca^2+^ signals (8, 11). Release of ALG-2 from either of these inhibitory states appears to switch its activity, causing ALG-2 homodimers to stimulate COPII targeting and ER export; this switch can be induced by short-lived–as opposed to continuous–Ca^2+^ signals (8). To summarize, it appears that Ca^2+^ signals, depending upon strength and duration, can induce either stimulatory or inhibitory actions of ALG-2 at ERES, and this regulation involves peflin, sec31A, and potentially other effectors.

But what governs ALG-2 influences at steady state? ER Ca^2+^ signals at steady state may be due to spontaneous Ca^2+^ oscillations mediated by gated Ca^2+^ channels, e.g. inositol trisphosphate receptors (IP3Rs) (12), which have been shown to dynamically alter ALG-2 localization (6), or else by continuous ER leak, for example mediated by presenilin-1 (13, 14) and/or the translocon (15, 16). A recent study in goblet cells, another polarized epithelial cell type, found that spontaneous cytosolic Ca^2+^ oscillations requiring ryanodine receptor (RyR) Ca^2+^ channels provide a steady-state signal acting as a tonic brake to mucin granule exocytosis (17). In this case, however, the Ca^2+^ sensor was apparently KChIP3 localized to pre-exocytic secretory granules. Much remains to be learned about steady-state ER Ca^2+^ signals and their effectors throughout the secretory pathway.

Since IP3 receptors, during agonist-driven Ca^2+^ signaling, contribute to PEF protein-mediated regulation of ER export, we wondered whether spontaneous Ca^2+^ signals regulated by IP3Rs at steady state might also contribute to setting the basal ER export rate. Here we demonstrate that normal rat kidney (NRK) epithelial cells participate in intercellular Ca^2+^waves (ICWs) that move back and forth across the monolayer under normal growth conditions. Depletion of IP3R-3, the major IP3R isoform in NRK cells, potentiates the remaining IP3Rs, causing greater participation in Ca^2+^ oscillations and ICWs in the absence of added agonists. Furthermore, this increased activity leads to more ALG-2 and Sec31A and less peflin at ERES, and increases the basal ER export rate of the COPII cargo VSV-G_ts045_ by up to 60%. Recruitment of more Sec31A to ERES by ALG-2 was client cargo-dependent suggesting that ALG-2 may cooperate with the sorting function of COPII. Indeed, more detailed studies indicated that the increased ER secretion was driven by greater capture and concentration of COPII client cargo at ERES at the expense of bulk flow cargo.

## RESULTS

### IP3R-3 depletion caused potentiation of agonist-dependent Ca^2+^ signaling

To perturb the pattern of Ca^2+^ oscillations under steady-state growth conditions, we focused on depletion of the IP3R-3 isoform. Previous literature indicated that the IP3R-3 isoform is anti-oscillatory in HeLa, COS, and NRK fibroblast cell lines (12, 18, 19). However, some literature suggests that decreased overall IP3R channel density should decrease basal signaling (18). Thus, empirical documentation of effects on Ca^2+^ handling in NRK epithelial cells (ATCC CRL-6509) was essential. To characterize the expression of IP3R isoforms in NRK cells, immunoblotting was performed using antibodies to all three individual IP3R isoforms - IP3R-1, IP3R-2, and IP3R-3. Both IP3R-1 and IP3R-3 were readily detectable in NRK extracts, but IP3R-2 was undetectable (data not shown). To determine the ratio of IP3R-1 and IP3R-3 in NRK cells, qRT-PCR was carried out on NRK cell total RNA using isoform-specific primers. As shown in Figure 1A, IP3R-3 was the major isoform, representing 67% of all IP3R subunits; the ratio of IP3R-3:IP3R-1 was thus 2:1 in NRK cells. Using siRNA, IP3R-3 could be depleted by 77% (Figure 1B). Though there was detectable co-depletion of IP3R-1, the ratio of R-1:R-3 isoforms was dramatically shifted toward R-1. The IP3Rs form both functional homo-tetramers and functional hetero-tetramers (12, 20). Hence, IP3R-3 depletion would eliminate most IP3R-3 homo-tetramers and IP3R-3-containing hetero-tetramers, leaving mostly IP3R-1 homo-tetramers, while constituting about one-third of the normal number of channels.

**Figure 1.**
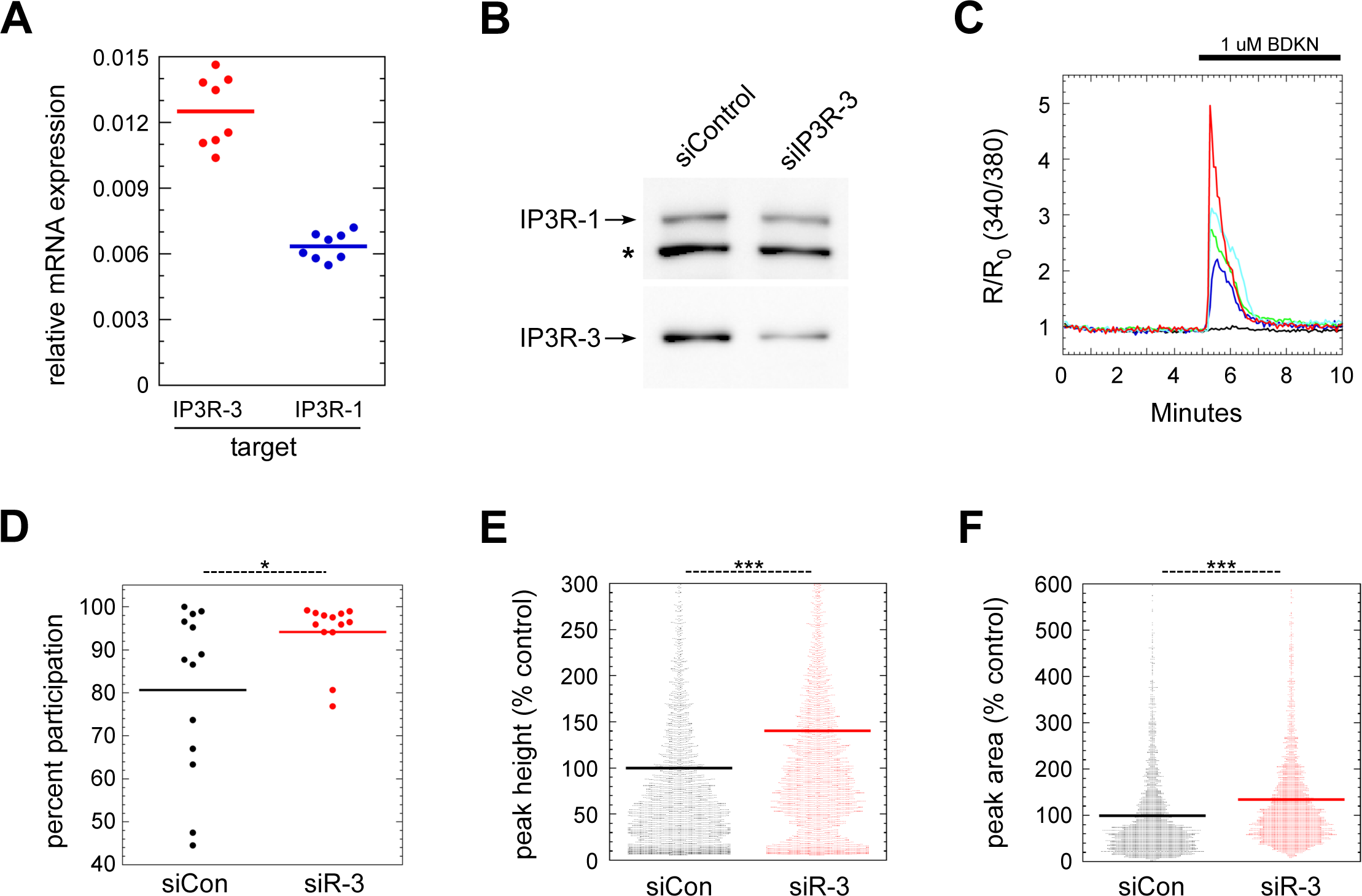
IP3R-3 depletion potentiates agonist-dependent Ca^2+^signaling. **(A)** Expression ratio of IP3R-1 and IP3R-3 in NRK cells analyzed by qRT-PCR. The quantity of each target RNA is normalized to that determined for GAPDH in parallel reactions. **(B)** Verification by immunoblot of IP3R-3 depletion using siRNA. Asterisk marks an unknown nonspecific band recognized by the anti-IP3R-1 antibody. **(C)** FURA-2 cytosolic Ca^2+^ recording of 5 control NRK cells in one field upon stimulation with 1 µM extracellular bradykinin (BDKN). Four cells displayed cytosolic Ca^2+^ signals of varying sizes while one (black trace) remained inactive. **(D)** Quantitation of the percentage of cells per field responding to BDKN stimulation as in (C). Each field contained a total of ∼300 cells. **(E,F)** Quantitation of peak height (defined as change from average baseline value) (E) and peak area (F) from traces of BDKN-challenged cells. Results in (E) contain all cells while (F) includes only the BDKN-responsive cells. Results in (D-F) consist of three replicate experiments combined. Cells were defined as BDKN-responsive in (D) and (F) if their peak height was ≥0.1. In (E,F), each dot represents one cell (N≈4000 cells per condition). p-values for two-tailed Student T-tests with unequal variance are indicated above; * = p ≤ .05; ** = p ≤ .005; *** = p ≤ .0005.

To indicate whether IP3R-3 depletion led to more promiscuous cytosolic Ca^2+^signaling, we first tested whether the remaining IP3R channels displayed potentiated responses to IP3-generating agonists. As demonstrated in Figure 1C, which displays representative FURA cell traces, most NRK cells responded to 1 µM bradykinin with a single synchronous surge of Ca^2+^ followed by a fall-off in Ca^2+^ during the ensuing 2-3 minutes (colored traces). About 20% of cells did not visibly respond (black trace), and cells with multiple, distinct oscillations were infrequently seen (not shown). Figure 1D-F displays results of systematic comparisons of control and IP3R-3-depleted cells using the bradykinin protocol from part C. Notably, as shown in Figure 1D, a significantly greater percentage of IP3R-3-depleted cells responded to bradykinin, raising the participation rate from ∼80 % to ∼95 %. Additionally, as shown in Figure 1E, IP3R-3-depleted cells responded with significantly higher amplitude Ca^2+^ surges than control cells, represented by a ∼40 % increase in peak amplitude. We also integrated the entire Ca^2+^ response, from onset of signaling until returning to baseline, among responding cells only, which demonstrated a highly significant ∼20 % increase in Ca^2+^ release (Figure 1F). In summary, Figure 1 C-F indicates that IP3R-3 depletion caused increased Ca^2+^ signals during the synchronous response to high doses of an IP3-generating agonist.

### IP3R depletion caused increased spontaneous Ca^2+^ oscillations and participation in intercellular Ca^2+^ waves

To characterize basal Ca^2+^ signals in unstimulated NRK cells, we monitored cytoplasmic Ca^2+^ in NRK cells using the Ca^2+^-sensitive dye FURA-2AM; cells were recorded for 20 min at 37 °C in regular growth medium containing 10% FBS in the absence of added agonists. As illustrated by representative individual cell traces in Figure 2A, some cells displayed a short burst of Ca^2+^ oscillations during the 20-minute interval. We created individual traces for all 514 cells in the image field, and determined that 52 cells or 10.1% displayed at least one burst of signaling; each burst contained 1-6 distinct oscillations in which each oscillation persisted for about 1 minute. Though it was not technically feasible to monitor for longer periods, we assume that over longer time intervals the majority of NRK cells would display similar episodes of spontaneous Ca^2+^ signaling.

**Figure 2.**
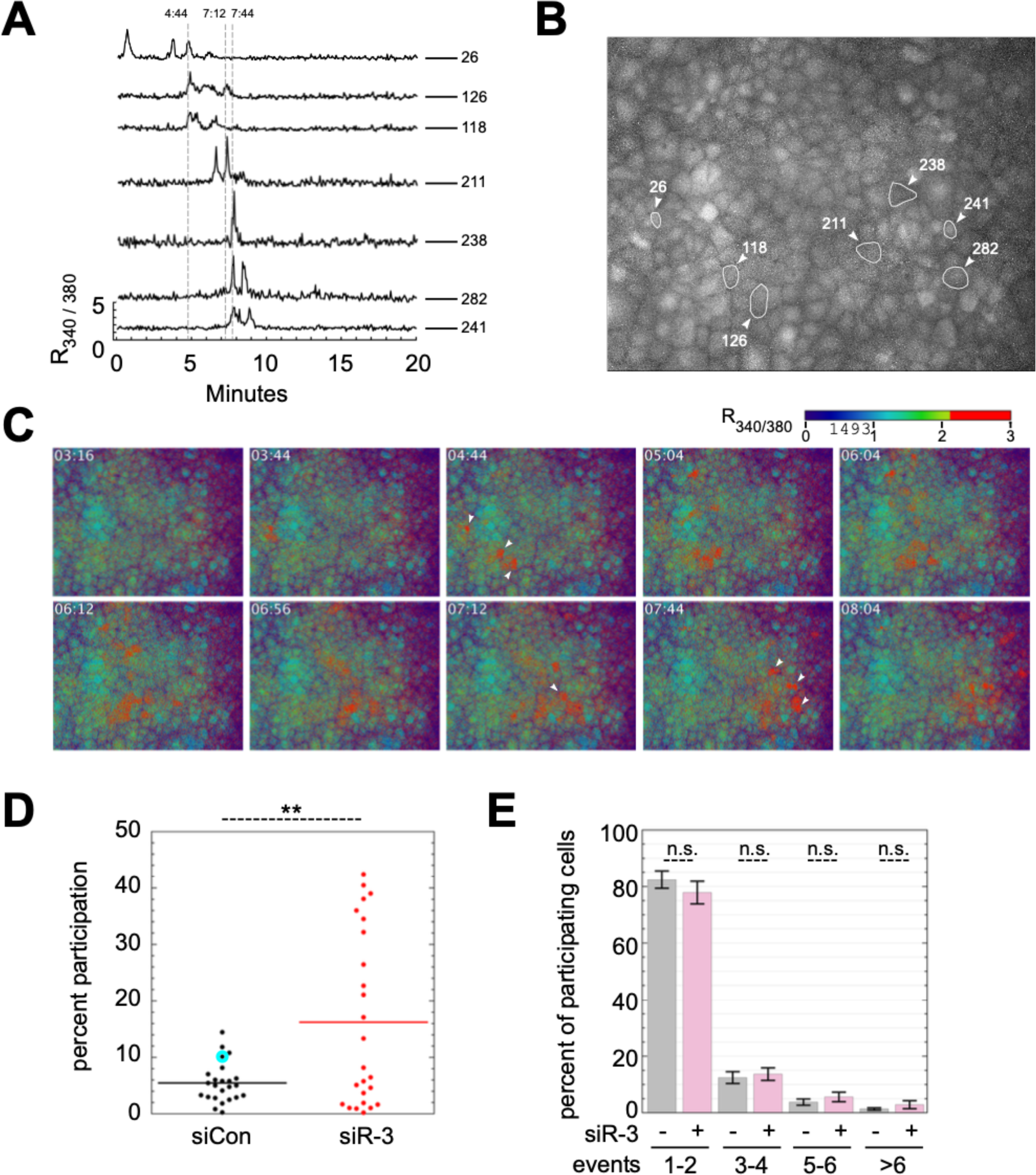
IP3R-3 depletion caused increased spontaneous Ca^2+^ oscillations. **(A)** Individual FURA-2 cytosolic Ca^2+^ traces of 7 control NRK cells in one field. **(B)** Group of 338 cells containing the same cells plotted in A, with their individual locations marked. **(C)** R_340/380_ heatmap of field in (B) at distinct timepoints. White arrows indicate position and time of major cytosolic Ca^2+^ signals in cells from (A). **(D)** Quantitation of cells participating in spontaneous cytosolic Ca^2+^ signals during the 20-minute imaging period as in (A), with each dot summarizing a run of ∼440 cells (49 individual runs, N=∼23,000 cells). The particular run shown in (A-C) is highlighted in cyan. **(E)** Quantitation of number of oscillations per cell from (D), as percent of all cells with oscillations. p-values for two-tailed Student T-tests with unequal variance are indicated above; * = p ≤ .05; ** = p ≤ .005; *** = p ≤ .0005. Error bars show SEM.

Intercellular coordination of Ca^2+^ signaling was typically observed in these recordings. Coordination took the form of groups of contiguous or nearby cells that oscillated almost precisely in unison. Furthermore, in most cases successive groups of activated cells would spread laterally over time, creating waves of high Ca^2+^ that traversed the monolayer. Figure 2B shows a group of 338 cells containing the same cells plotted in Figure 2A, with their individual locations marked. Figure 2C displays the same group of cells at several distinct timepoints during the recording with the R_340/380_ ratio displayed using a heatmap for contrast. In detail, cell 26 initiated a major Ca^2+^ wave at t = 04:44, when cells 118 and 126 joined and oscillated in unison a few seconds behind cell 26. Five to ten cells at a time then oscillated together in successive groups that led upward and to the right across the field, culminating in cells 238, 241, 282 and others oscillating in unison at t = 7:44 before the wave moved further rightward and out of view. The continuous nature and high local participation in this intercellular Ca^2+^ wave (ICW) is demonstrated in the video (Supplementary Video File 1) containing all 150 timepoints during the first 10 minutes of the recording. The mechanism and functions of ICWs are not well understood; they can be driven by both cell-to-cell diffusion of Ca^2+^ and IP3 through gap junctions and also by paracrine purinergic signaling (PPS) involving vesicular and non-vesicular release of the Ca^2+^ agonist ATP (21). While we did not investigate the mechanisms of NRK cell ICWs further, we speculate that one relevant consequence of ICWs is that they may help amplify and normalize cell participation in spontaneous, steady-state Ca^2+^ signaling.

We compared control and IP3R-3-depleted NRK cells for their participation in spontaneous, steady-state Ca^2+^ signaling using recordings similar to that described above. Each of the 300-500 cells in the view area of each run was analyzed using a semi-automated workflow in ImageJ and R that generated individual Ca^2+^ traces that could be rapidly scrolled through for manual scoring; each trace was scored for the number of spontaneous Ca^2+^ oscillations, from which the percentage of cells participating in oscillations was derived for each run. As shown in Figure 2D, which represents scoring of ∼23,000 cells in 49 individual runs, IP3R-3-depleted cells exhibited significantly higher cell participation in spontaneous signaling, increasing about three-fold, from 5.5% to 16% cell participation during each 20-minute period. Though it was not possible to objectively score ICWs *per se*, a majority of recordings displayed groups of cells oscillating synchronously, indicating that intercellular coordination or spread of Ca^2+^ oscillations may be a major driver of participation in basal Ca^2+^ signaling in NRK cells. The specific recording shown in Figure 2A-C is highlighted in the composite plot (2D) with a cyan halo.

While this recording displayed the linear cell-to-cell spread of signaling unusually clearly, it was not remarkably active in terms of cell participation, which ranged much higher among IP3R-3-depleted cells. We also compared the number of oscillations per cell among participating cells between control and IP3R-3-depleted cells, and found that there was a trend toward greater numbers of oscillations in IP3R-3-depleted cells (Figure 2E). In conclusion, IP3R depletion caused increased spontaneous Ca^2+^ oscillations, facilitated through ICWs.

### Effects of IP3R-3 depletion on ER Ca^2+^ stores

We also characterized ER luminal Ca^2+^ stores in control and IP3R-depleted cells using D1ER, a genetically encoded ER luminal FRET-based Ca^2+^ sensor (22). As shown by representative traces in Figure S1A, upon addition of ionomycin and EGTA to live NRK cells, R/R_0,_ which for the first five minutes has a value near 1, quickly plunged and approached an R_min_/R_0_ value; R_min_/R_0_ ranged from ∼0.75 (for control cells) to ∼0.85 (for IP3R-depleted cells). The difference between the initial R/R_0_ and R_min_/R_0_–or ΔR/R_0_–was interpreted to represent the resting ER luminal Ca^2+^ concentration of the cell. As shown in Figure S1B, which combines cells from multiple replicate experiments, ΔR/R_0_ was 28 % smaller in IP3R-depleted cells, indicating a lower resting ER Ca^2+^ concentration. To verify the conclusion using a different approach, we used an irreversible sarco-endoplasmic reticulum Ca^2+^ ATPase (SERCA) inhibitor, 1 µM thapsigargin (Tg), to release all ER Ca^2+^, and then measured the magnitude of the resulting cytoplasmic Ca^2+^ surge in NRK cells transfected with the genetically encoded FRET-based cytosolic Ca^2+^ sensor D3cpv (23). As shown in Figure S1C and D, IP3R-3-depleted NRK cells displayed smaller Ca^2+^ releases, by 33%, upon Tg addition, consistent with the conclusion from direct luminal measurements (parts A and B) that ER Ca^2+^ stores were ∼30% reduced in IP3R-3-depleted cells. We also used a similar Tg addition protocol together with a genetically encoded FRET-based mitochondrial matrix Ca^2+^ sensor, 4mtD3cpv (24), and found that mitochondria took up less Ca^2+^ upon release of ER Ca^2+^ in IP3R-depleted cells (data not shown). Together these experiments indicate that, for reasons that are not obvious, depletion of IP3R-3 in NRK cells results in partial reduction of ER Ca^2+^ stores. We do not believe the increased spontaneous signaling was responsible, though we cannot eliminate that mechanism. Further speculations on the molecular cause of the Ca^2+^ store reduction are offered in the Discussion.

### IP3R depletion down-regulates PEF protein expression but does not alter expression of multiple other trafficking machineries

To begin the examination of IP3R-3 depletion effects on ERES and ER-to-Golgi transport, we asked whether IP3R depletion changed expression of trafficking machinery proteins. In particular, one could imagine that altered ER Ca^2+^ handling could result in unfolded protein response (UPR) activation which is known to increase expression of COPII-related components. We first examined relevant gene expression using qRT-PCR. As shown in Figure 3A, IP3R-3 knockdown did not significantly affect mRNA expression of COPII inner coat components Sec24A, B, C, or D, nor outer shell component Sec31A, nor ERES scaffolds Sec16A and B. There was perhaps a trend toward an increase of Sec24C, but it was not statistically significant and not supported by immunoblotting of cell extracts (see below). Overall, the qRT-PCR data did not support general up-regulation of ERES components as would be expected if basal UPR signaling were higher. In addition, we found that none of the UPR-related genes ATF6, CHOP, or Grp78 were affected by IP3R depletion. We found that ALG-2 and peflin were expressed at very similar levels and not affected by IP3R depletion at the transcript level. Finally, Bcl-2 transcripts were reduced by a highly significant ∼30% upon IP3R-3 depletion; this demonstrates that IP3R-3 depletion influenced a gene product known to directly interact with and regulate the activity of IP3Rs (25). We complimented this analysis with immunoblotting of NRK cell extracts transfected with control or IP3R-3 siRNAs. As seen in Figure 3B, UPR mediators phospho-EIF2α, phospho-IRE-1 and ATF4 did not noticeably change upon IP3R depletion. Turning to the transport machinery, Figure 3C demonstrates no changes in coat subunits Sec31A, ß-COP, Sec24c, Sec23, the ER/Golgi tether p115, cargo receptors ERGIC-58 (labeled “p58”) and p24. These imply that IP3R depletion did not alter transport by changing expression of the required transport machinery. However, we also observed in immunoblots a small yet consistent decrease in both ALG-2 and peflin. This effect was quantified from multiple biological and technical replicates and displayed in Figure 3D, revealing on average a ∼25% decrease in both proteins. Since mRNA levels were not affected (Figure 3A), it appears that this change occurs via a post-transcriptional mechanism. Note that concomitant reduction of both total cellular ALG-2 and peflin is not predicted to in and of itself significantly change ER-to-Golgi transport, since the two are still present in the same ratio. However, both proteins are high turnover proteins that are destabilized when not in the ALG-2-peflin heterodimer (10) which is disrupted by elevated Ca^2+^ (9). Hence, their reduced expression might reflect that the more dynamic cytosolic Ca^2+^ environment in the IP3R knockdown condition creates greater flux between the ALG-2-containing complexes.

**Figure 3.**
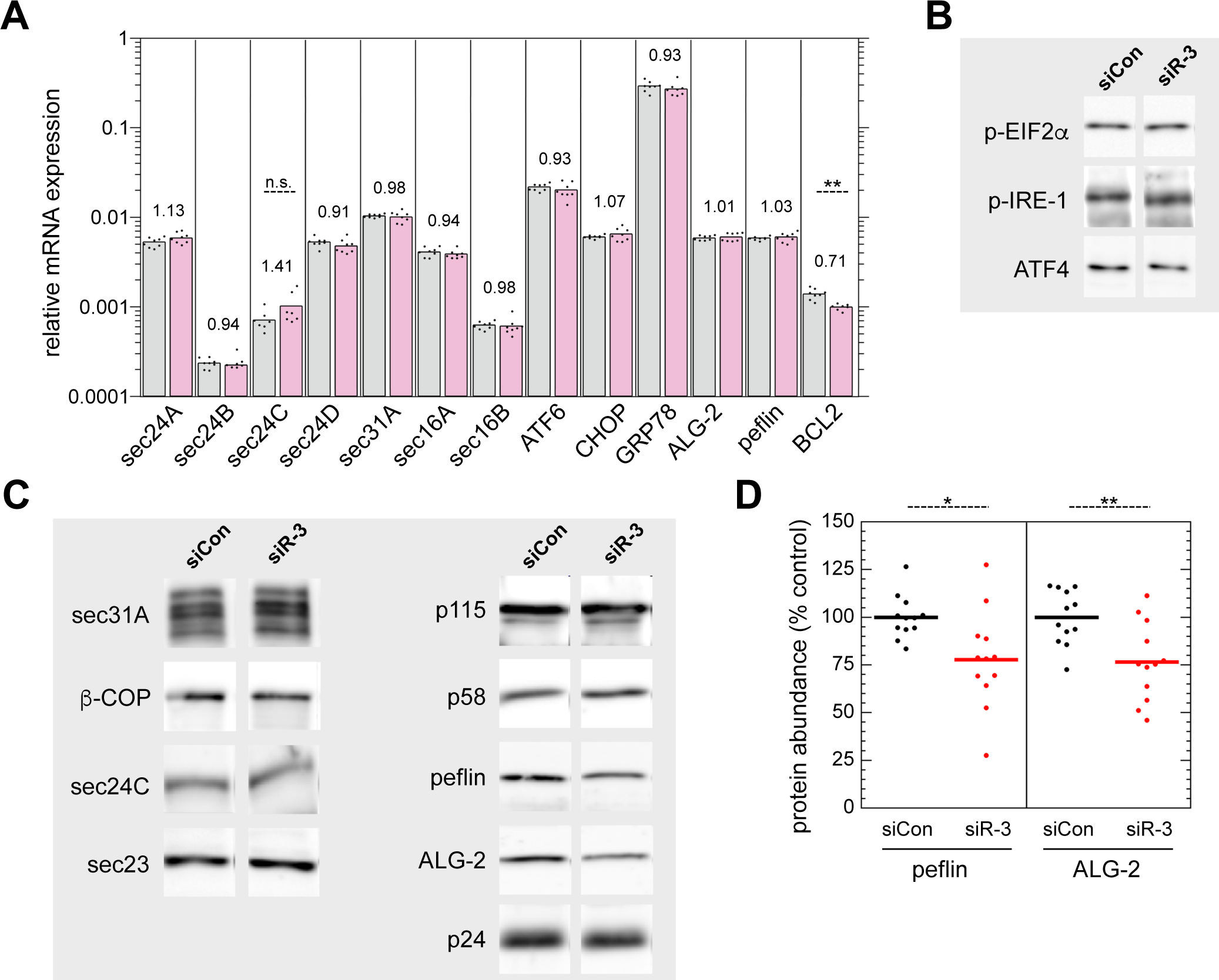
IP3R-3 depletion destabilizes ALG-2 and peflin but does not affect expression of other ER export machinery. **(A)** NRK cells were transfected with control (gray) or IP3R-3 (light red) siRNAs, grown for 48 h, lysed, and then mRNA was analyzed by qRT-PCR to detect expression of trafficking and other proteins. The quantity of each target RNA is normalized to that determined for GAPDH in parallel reactions. Value labels indicate the ratio of expression in siR-3 cells to that of control cells. **(B, C)** Immunoblots of ER stress proteins (B) and ER/Golgi trafficking machinery (C) in NRK cell extracts, 48 h after transfection with control or IP3R-3 siRNAs. No significant changes were observed, except for decreases in ALG-2 and peflin. **(D)** Quantitation of multiple technical replicates from several different experiments documented effects of IP3R-3 depletion on ALG-2 and peflin protein abundance. p-values for two-tailed Student T-tests with unequal variance are indicated above; * = p ≤ .05; ** = p ≤ .005; *** = p ≤ .0005.

### IP3R depletion causes increased ALG-2 and Sec31A, but decreased peflin targeting to ERES

The mild destabilization of ALG-2 complexes during IP3R-3 depletion (Figure 3D) led us to ask whether ALG-2 and peflin targeting to ERES was altered, since this could alter COPII coat targeting and thus ER export. ALG-2 binds Sec31A at ERES either as a peflin-ALG-2 heterodimer, or as the ALG-2 homodimer. While the homodimer is generally stimulatory, the ALG-2-peflin heterodimer is inhibitory (8). We carried out immunofluorescence of endogenous ALG-2, peflin, and Sec31A on control and IP3R-3-depleted NRK cells. For a non-COPII-related marker of ERES we used rBet1, an ER/Golgi SNARE whose distribution is mostly restricted to ERES and ERGIC elements (26). Representative images of cells are shown in Figure 4A-C. We segmented the labeling patterns into discrete objects by thresholding and then used Boolean image math to measure the areas of objects and areas of co-localization between objects representing different proteins in the same cell (8, 11). As shown in Figure 4D, when IP3R-3 was depleted, co-localization between peflin and rBet1 objects decreased by ∼25%. The decreased co-localization was comprised of both a decrease in the percentage of total peflin objects that co-localized with rBet1, and a decrease in the percentage of total rBet1 objects that co-localized with peflin (data not shown); this reciprocity of effects, which was also true in parts E and F, establishes a true decrease in co-localization that could not result from mere changes in labeling intensity or expression. At the same time, co-localization between ALG-2 and rBet1 significantly increased by ∼35% in IP3R-3-depleted cells (Figure 4E). Focusing on the outer coat, we found that the co-localization area of Sec31A and rBet1 increased by ∼50% upon IP3R-3 depletion (Figure 4F). The expansion of Sec31A-rBet1 overlap most likely indicates that more outer coat was recruited to ERES, consistent with past work concluding that ALG-2 ‘stabilized’ the outer coat at ERES (4, 7). Taken together the immunofluorescence changes during IP3R depletion– decreased peflin, increased ALG-2, and increased Sec31A recruitment to ERES–indicate that the increased steady-state Ca^2+^ signaling was sufficient to trigger changes to ALG-2 targeting in a pattern shown previously to increase ER-to-Golgi transport rates (8, 11).

**Figure 4.**
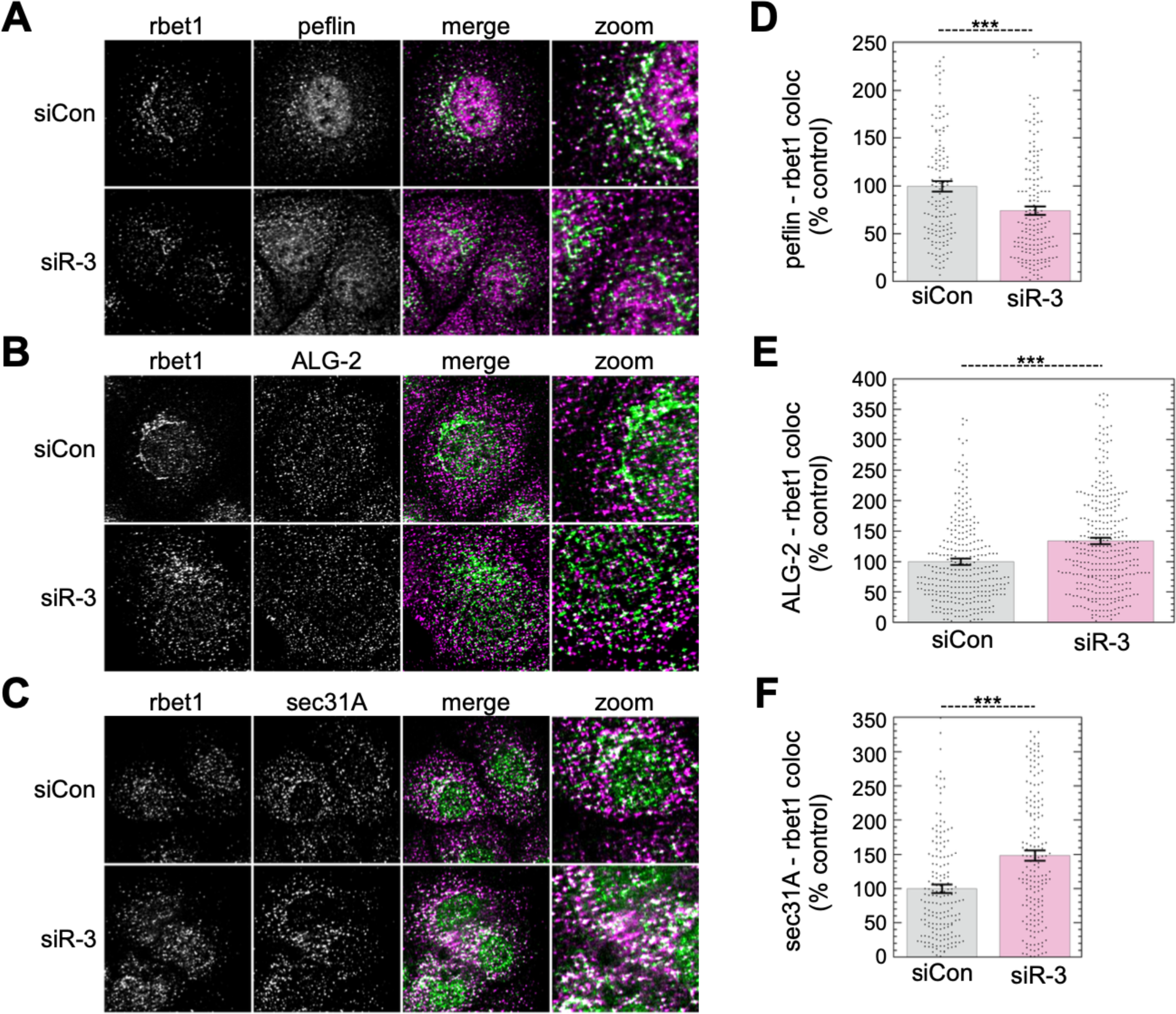
IP3R depletion causes a decrease in peflin and increases in ALG-2 and sec31A at ERES. **(A-C)** NRK cells were transfected with the indicated siRNAs. After 48 h, cells were fixed and immunolabeled for endogenous rbet1, peflin, ALG-2, or sec31A, as indicated. Representative micrographs are shown. **(D-F)** Object analysis on ∼150 cells (D, F) or ≥300 cells (E) per condition to quantify overlap area between rbet1 objects and the other labeled proteins. p-values for two-tailed Student T-tests with unequal variance are indicated above; * = p ≤ .05; ** = p ≤ .005; *** = p ≤ .0005. Error bars show SEM.

### Depletion of IP3R-3 adjusts ER-to-Golgi transport in a cargo-specific manner

To test the effects of IP3R-3 depletion on ER-to-Golgi cargo transport, we used an intact-cell transport assay (2, 8, 11, 27) employing three different synchronized release mechanisms for the transmembrane COPII client cargo VSV-G as well as a synchronized bulk-flow cargo. Cargo was released from the ER for 10 minutes, after which the cells were fixed, and a transport index was calculated consisting of the intensity of the cargo in the Golgi divided by the intensity remaining in the ER. For the GFP-tagged VSV-G constructs, we employed: 1) temperature-sensitive GFP-VSV-G_ts045_ which traffics following shift from 41 to 32 °C; 2) GFP-FM_4_-VSVG_tm_, which contains a luminal GFP and controlled aggregation domain connected to the wildtype VSV-G transmembrane domain and cytoplasmic C-terminus (8). Here, addition of the highly specific ligand D/D Solubilizer causes dissociation of large cargo assemblies allowing packaging into COPII vesicles (28, 29). 3) RUSH-VSV-G-GFP (30, 31), which traffics after added biotin outcompetes binding of the construct to an ER-localized streptavidin “hook”. For the bulk-flow cargo, we employed GFP-FM_4_-hGH, an entirely luminal construct containing the controlled aggregation domain and hGH and no known functional sorting motif. Figure 5A shows example images and illustrates the morphological principle of the assay. Importantly, as demonstrated quantitatively in Figure 5B, knockdown of IP3R-3 resulted in up to a 60% acceleration of the ER-to-Golgi transport rate of all three VSV-G constructs. In parallel, IP3R-3 depletion decreased trafficking of the bulk flow construct by ∼25%. The use of three distinct VSV-G constructs eliminates the possibility that structural rearrangements required for any particular synchronization method, for example folding or disaggregation kinetics, are required to generate the transport differences. The cargo specificity of the effects (GFP-FM_4_-VSV-G_tm_ vs. GFP-FM_4_-hGH) is identical to that caused by direct activation of ALG-2 in basal Ca^2+^ through siRNA depletion of its inhibitory subunit peflin (8), and thus supports that ALG-2 drives the phenomenon. While the cargo specificity could be indicative of changes in COPII cargo sorting, it could also be indicative of distinct trafficking pathways.

**Figure 5.**
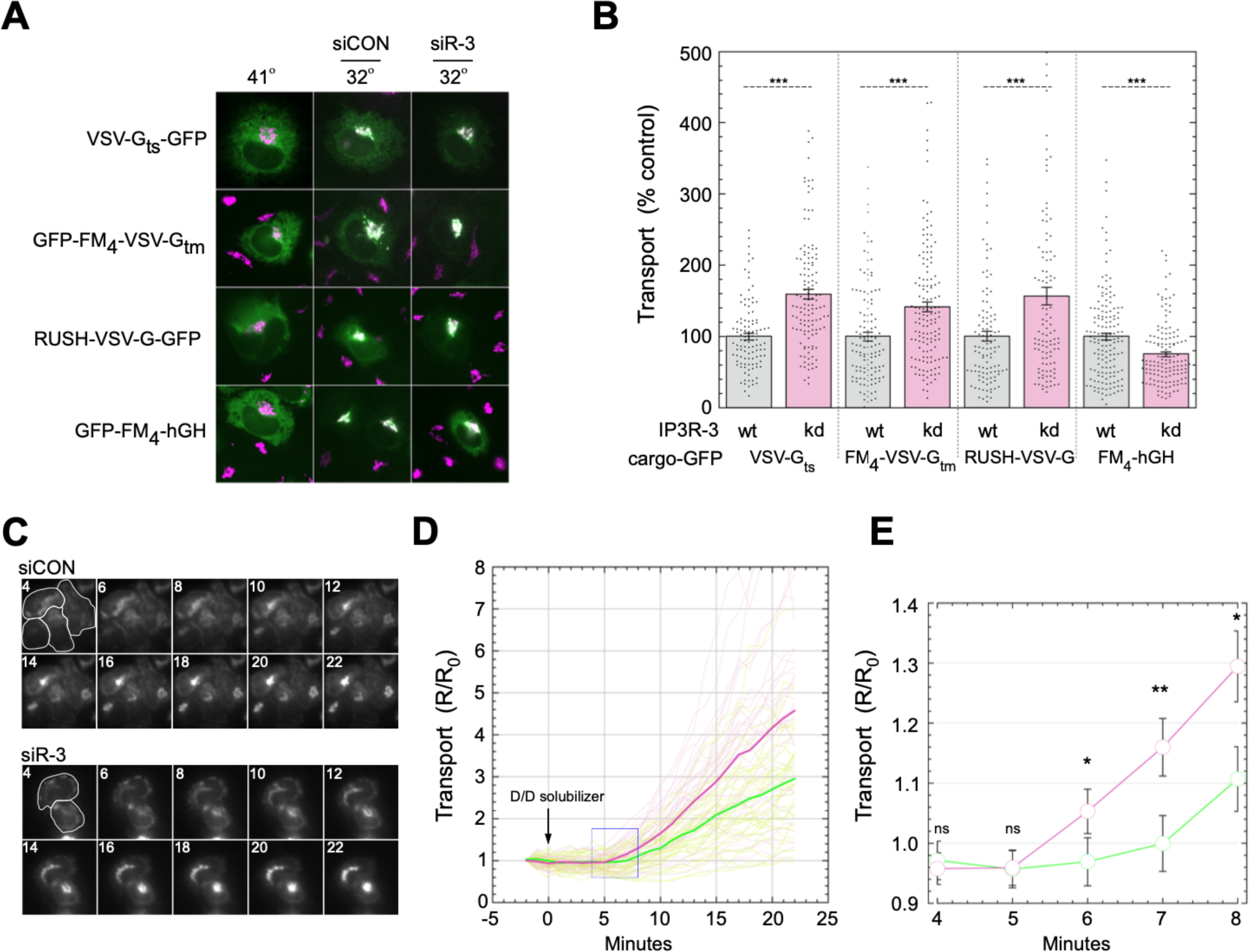
Expression of IP3R-3 regulates ER-to-Golgi transport in unstimulated NRK cells. **(A)** Representative images of NRK cells from the ER-to-Golgi transport assay quantified in B. NRK cells were transfected with VSV-G_ts045_-GFP with or without siRNAs as indicated. Following growth at 41 °C for 48 h, cells were shifted to 32 °C for 10 min, to permit transport, prior to fixation. Fixed cells were immuno-labeled with mannosidase II and imaged by widefield microscopy; VSV-G-GFP and mannosidase II channels are displayed for each cell. **(B)** Each transfected cell was assigned a transport index representing trafficking of VSV-G based upon the ratio of Golgi to peripheral green fluorescence. The transport index of each individual cell is plotted. 200-300 cells were randomly quantified from each condition, and results shown are representative of at least three experiments with consistent trends. **(C-E)** A live-cell transport assay employing control and IP3R-3 siRNAs and the FM_4_-VSV-G_tm_ cargo. **(C)** Representative cells imaged in the GFP channel; cell outlines are show on the first panel and minutes post addition of D/D solubilizer are indicated on each. **(D)** R/R_0_ traces of NRK cell transport indexes from a live-cell ER-to-Golgi transport assay. Individual cell traces are shown as faint lines while average traces are shown in bold. Control and siIP3R-3 are shown as green and red, respectively. Transport index as in (B) was calculated at each timepoint and expressed as a ratio to the initial value for each cell. N=40 cells per condition. **(E)** Closer view of average traces from (D) showing first timepoints with measurable ER-to-Golgi transport. p-values for two-tailed Student T-tests with unequal variance are indicated above; * = p ≤ .05; ** = p ≤ .005; *** = p ≤ .0005. Error bars show SEM.

We also tested the effects in a live-cell assay which generated a kinetic transport timecourse for each cell. In this experiment, we employed GFP-FM_4_-VSV-G_tm_. At 37 °C 4 minutes after addition of D/D solubilizer, the cargo was restricted to the ER in both control and IP3R-3-depleted cells (Figure 5C, “4”). Cargo first became visible in the Golgi 6-7 minutes following ligand addition and accumulated more or less linearly for the subsequent 15 minutes. Upon quantitation of 40 cells per condition from sequential runs, it became clear that the IP3R-depleted population contained many more high-transporting cells, while the control cells contained many more low-transporting cells (Figure 5D). Furthermore, upon close examination of the timecourses at t=4 to t=10 m, it became clear that transport in the control cells exhibited a longer lag between ligand addition and the linear accumulation phase of cargo in the Golgi (Figure 5E). While the eventual slopes of the transport curves were also different, the pronounced difference at the onset of transport implies that the secretory machinery functioned at a higher level from the start of the reaction. Direct effects on an early step in transport such as cargo sorting or ER export are fully compatible with the ER-to-Golgi kinetics we observed.

### The increased transport of VSV-G requires ALG-2 and increased cytosolic Ca^2+^ signals

Since ALG-2 binds Sec31A in response to Ca^2+^ signaling (5, 6), we tested whether ALG-2 is required for the increased transport observed in IP3R-3-depleted cells. As shown in Figure 6A and quantified in Figure 6B, IP3R-3 depletion in otherwise wildtype cells increased transport by ∼60% (bars 1 vs. 2), consistent with Figure 5. However, in ALG-2-depleted cells, depletion of IP3R-3 caused no further significant effect (bars 3 vs. 4, p=.257). This is the result expected if ALG-2 is required to mediate the increased transport due to IP3R depletion; ALG-2 is epistatic to IP3R-3 for transport effects. Loss of ALG-2 itself is slightly stimulatory in otherwise normal cells (bars 1 vs. 3), and slightly inhibitory in IP3R-3-depleted cells (bars 2 vs. 4). Hence it appears that the increased Ca^2+^ signaling in IP3R-depleted cells switched ALG-2 influence from that of a net inhibitor to that of a net stimulator of ER export.

**Figure 6.**
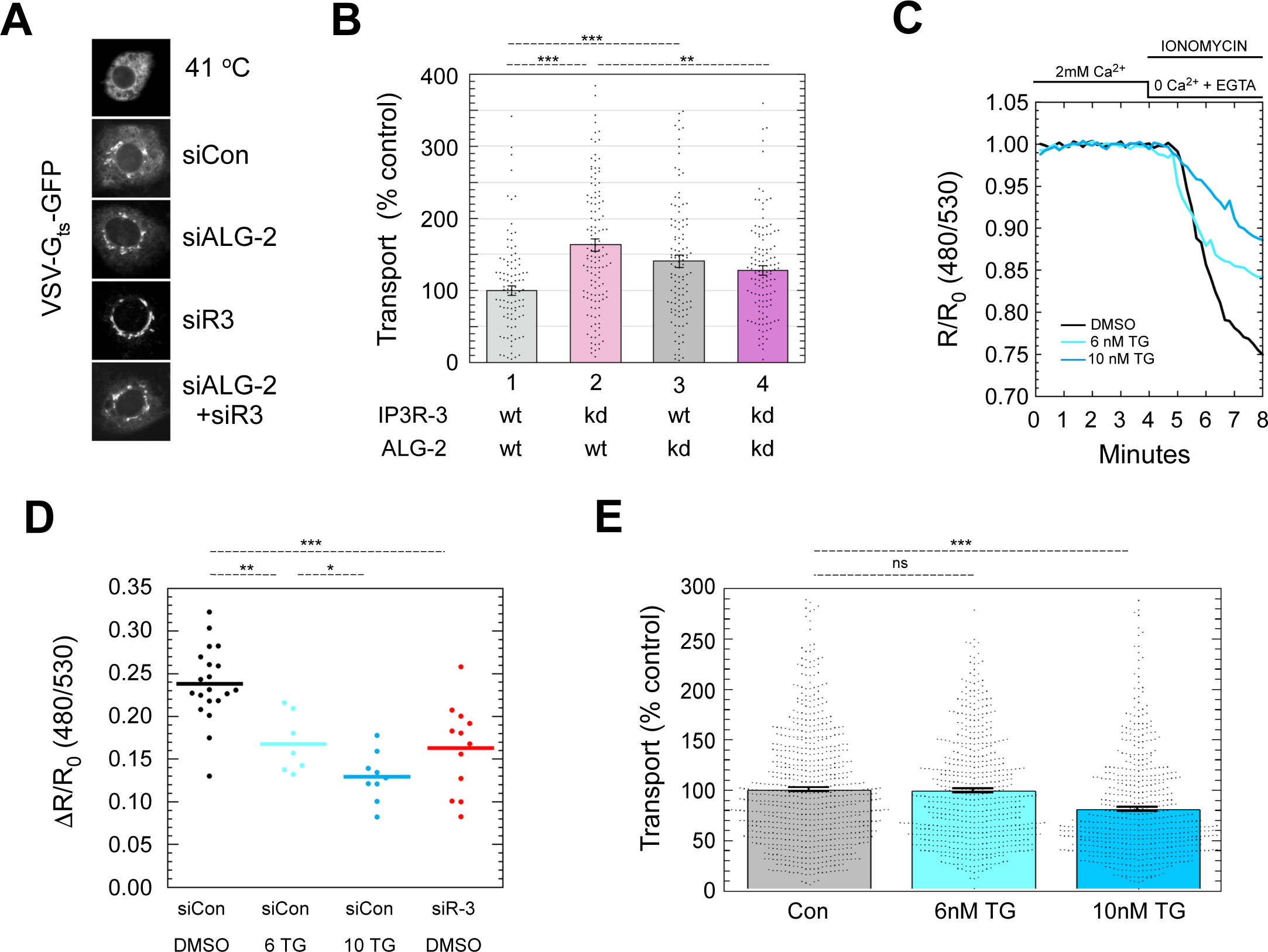
The increased transport of VSV-G requires ALG-2 and is not caused by decreased luminal Ca^2+^ stores. **(A)** Representative GFP images from a 10-min transport assay in NRK cells using VSV-G_ts_-GFP cargo and siRNA. **(B)** Quantitation of ER-to-Golgi transport as in Figure 5B. **(C)** Example luminal Ca^2+^ depletion traces from representative live cells using the ER FRET-based Ca^2+^sensor D1ER. Ca^2+^, ionomycin, and EGTA are added and withdrawn using continuous perfusion as indicated above. The emission ratio for CFP and YFP was determined using 430 nM illumination at each timepoint and expressed as a ratio to initial value. Cells were treated with the indicated TG concentration for 24 h prior to assay. **(D)** Drop in FRET ratio between 0 and 8 min was determined as in part C for several cells per run in multiple runs in cells treated with TG as in (C) or with IP3R-3 siRNA. **(E)** NRK cells were treated with the indicated concentrations of TG as in parts (C) and (D) and subjected to a standard 10-min ER-to-Golgi transport assay using VSV-G_ts_-GFP cargo. p-values for two-tailed Student T-tests with unequal variance are indicated above plots; * = p ≤ .05; ** = p ≤ .005; *** = p ≤ .0005. Error bars show SEM.

Since ALG-2 is a cytosolic Ca^2+^ sensor, the data so far imply that the increased frequency of cytosolic Ca^2+^oscillations upon IP3R-3 depletion causes ALG-2 to raise the basal secretion rate. However, the reduced luminal Ca^2+^ observed earlier (Figure S1) could present a challenge to the interpretation of effects on ER export, since it cannot be eliminated that a reduced luminal Ca^2+^store *per se* could somehow regulate the transport machinery by an unknown mechanism. If this were the case, comparably reduced luminal Ca^2+^ stores would still stimulate transport, even without depletion of IP3Rs and increased Ca^2+^ oscillations. We experimentally created such a condition by treating cells with very low levels of thapsigargin–two orders of magnitude below the concentrations typically used to induce UPR. As shown in Figure 6C and D, 6 nM Tg for 24 hours resulted in partial depletion of luminal Ca^2+^ similar to that caused by IP3R-3 siRNA, while at 10 nM, the Tg caused greater depletion. We used 24 hours to simulate any potential long-term effects on secretion as well as to get well beyond any stimulatory effects caused by the initial wave of released ER Ca^2+^. We then tested the effects of these partial ER Ca^2+^ depletions on ER-to-Golgi transport. As shown in Figure 6E, there were no significant effects on ER-to-Golgi transport at 6 nM Tg, but ∼20% inhibition of transport became apparent at 10 nM Tg. Inhibition of ER-to-Golgi transport under conditions of strong ER depletion is expected from our previous results (1, 2). The lack of transport effects at 6 nM demonstrates that partial luminal Ca^2+^ depletion for ∼24 hours does not cause the increased transport observed in IP3R-depleted cells. In conclusion, the data favor the interpretation that increased spontaneous cytosolic Ca^2+^ oscillations, and not the lowered Ca^2+^ store, primarily drives the ALG-2-dependent ERES changes that result in increased transport when IP3R-3 is depleted.

### ALG-2 activation produces higher concentrations of VSV-G at ERES

If ALG-2 increased sorting stringency for client cargo, one would expect client cargo to be more concentrated in ERES prior to export. Early in a transport reaction, it is difficult to quantify cargo specifically at ERES due to the very high level of cargo throughout the entire ER. In one alternative approach, we allowed cargo to accumulate significantly at ERES during a temperature block. At 10 °C, VSV-G_ts045_ can fold and be loaded into ERES in a coat-dependent manner, but vesicle budding and further steps in transport cannot occur (32). We activated ALG-2 Ca^2+^-dependently by IP3R-3 depletion or Ca^2+^-independently by depleting its inhibitory subunit peflin (8, 11), and then allowed VSV-G_ts_-GFP to accumulate for 1 hr at 10 °C. As shown in Figure 7A, this resulted in prominent accumulations of cargo in spots that mostly co-localized with Sec31A. Cargo was visualized using GFP fluorescence as well as immunofluorescence using the I14 conformation-sensitive monoclonal antibody specific for conformationally mature VSV-G trimers (33), which agreed well with GFP fluorescence. It appeared that cargo spots were brighter in IP3R-3-depleted and Peflin-depleted cells than in control cells, which was confirmed by quantitation of spot prominence. As shown in Figure 7B, spot prominence of GFP fluorescence was significantly increased by 100% in IP3R-3-depleted cells and 125% in Peflin-depleted cells. Spot prominence was defined as the maximum intensity of each spot divided by the mean intensity of a doughnut-shaped ROI immediately surrounding but excluding the spot. Total GFP fluorescence in the same set of cells was not significantly different (Figure 7C). Parallel standard ER-to-Golgi transport assays at the permissive temperature demonstrated qualitatively similar effects (Figure 7D), suggesting that concentration of cargo at ERES may account for the effect on overall transport. In sum, this experiment demonstrates that at 10 °C, activated ALG-2 drives higher concentrations of a client cargo at ERES.

**Figure 7.**
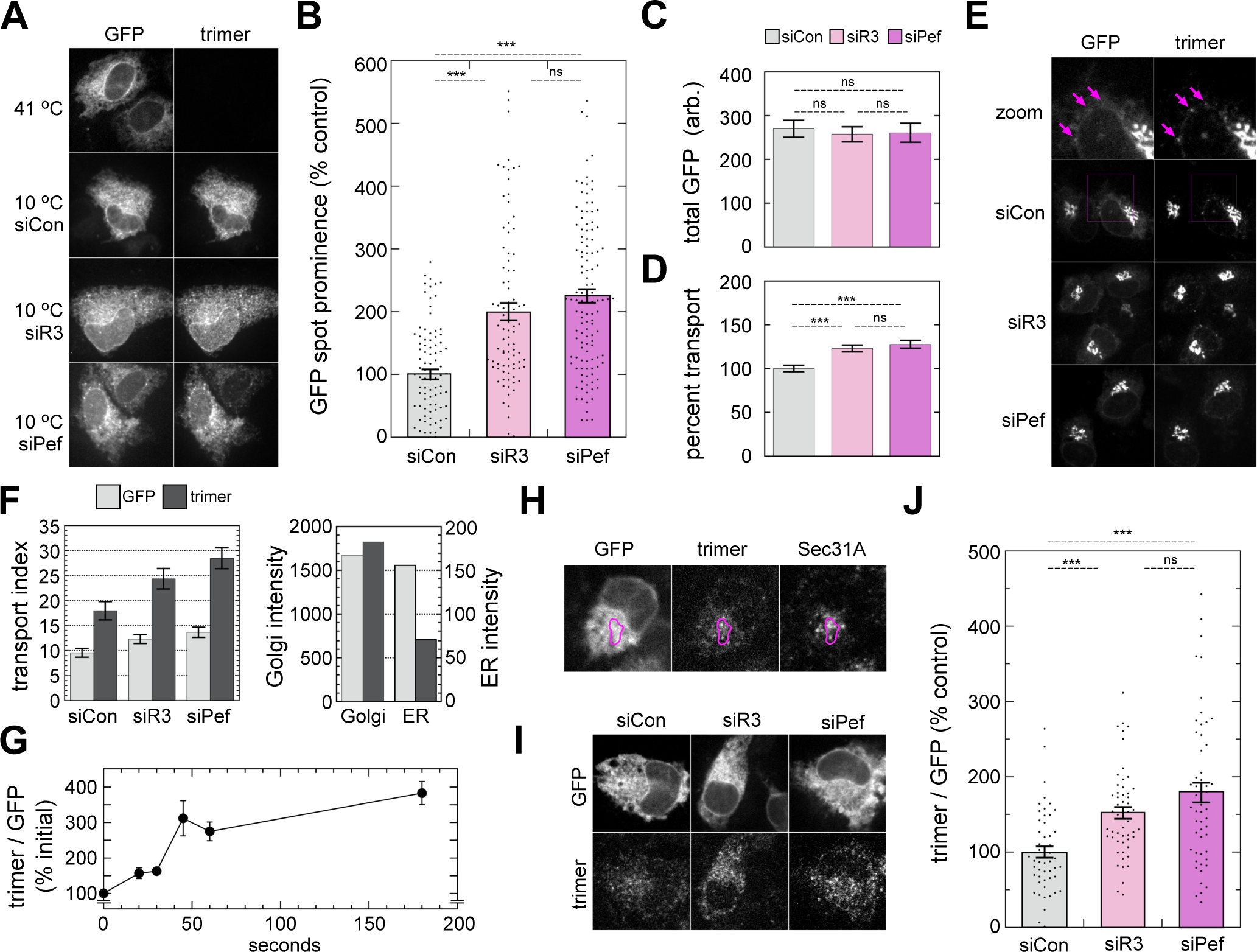
ALG-2 activation drives higher concentrations of VSV-G at ERES. **(A)** NRK cells transfected with the indicated siRNAs and VSV-G_ts_-GFP cargo were either kept at 41 °C or incubated at 10 °C for 1 hr prior to fixation and immunostaining with the I14 VSV-G conformation-specific antibody. Shown are the GFP channel (left) and trimer-specific channel (right) of the same cells. **(B)** Quantitation of GFP spot prominence in cells from (A). Spot prominence is defined as the maximum intensity of each spot divided by the mean intensity of the area immediately surrounding the spot (see methods). **(C)** Quantitation of GFP intensity of the entire cytoplasmic area (nucleus excluded) of the same cells quantified in (B). **(D)** Standard 10-min transport assay on parallel coverslips from the same experiment as (B). **(E)** 10-min ER-to-Golgi transport assay using the indicated siRNAs and VSV-G_ts_-GFP cargo. Images show GFP channel (left) and trimer-specific channel (right) of the same cells. Arrows indicate puncta prominent in the trimer channel that are weaker or absent in the GFP channel. **(F)** *Left,* quantitation of the transport assay from (E) with raw transport index values shown from the GFP and trimer channels. *Right*, individual components of the transport assay; Golgi intensity is plotted on the left y-axis while ER intensity is plotted in the right y-axis. Plotted values represent the average of all the siRNA conditions shown on the left plot. **(G)** Timecourse of appearance of trimer immunostaining upon switch from 41 °C to 32 °C. Trimer intensity of the entire cytoplasmic area was normalized the GFP intensity of the same area to counteract differences in transfection efficiency. **(H)** Colocalization at 60-sec of GFP, trimer, and Sec31A. ROI highlights area containing examples of precise (bottom) as well as imperfect (top) co-localization of trimer and Sec31A fluorescence. **(I)** 60-sec following shift to 32 °C, cells that had been transfected with the indicated siRNAs and VSV-G_ts_-GFP were fixed and immunolabeled with trimer-specific antibody. Representative cells are shown in the GFP (top) and trimer (bottom) channels. **(J)** Quantitation of cells from (I) using the same method as in (G). p-values for two-tailed Student T-tests with unequal variance are indicated above plots; * = p ≤ .05; ** = p ≤ .005; *** = p ≤ .0005. Error bars in all panels show SEM.

In a distinct approach that did not rely on a transport block, we returned to the conformation-specific anti-VSV-G monoclonal antibody, I14. Using immunoprecipitation with I14, we previously demonstrated that VSV-G_ts045_ in NRK cells completely folds and assembles into trimers within 3 minutes upon shift to the permissive temperature (33), while others have used more detailed velocity sedimentation approaches to reach similar kinetic conclusions (34). Here, we characterized I14 immunofluorescence properties during ER-to-Golgi transport. Using our standard ER-to-Golgi transport assay with VSV-G_ts045_-GFP at t=10 minutes, we found that GFP and trimer fluorescence did not match, unlike above during accumulation of trimer at 10 °C. Instead, during transport it appeared that trimer labeling lacked most of the general ER component and greatly emphasized cargo accumulations at ERES (Figure 7E, magenta arrows). When the transport assay was quantified separately in both channels (GFP and trimer), we found that both channels reported increased transport in cells depleted of IP3R-3 and in cells depleted of the ALG-2 inhibitory subunit peflin (Figure 7F, left plot). However, the trimer channel reported transport indices that were consistently about two-fold higher. Individual components comprising the transport index–Golgi and ER fluorescence–revealed that the higher transport indices reported by the trimer were indeed driven primarily by decreased detection of VSV-G in the general ER (Figure 7F, right plot). Since VSV-G_ts045_ completes folding within ∼3 minutes, it would be expected that I14 should detect the yet-to-be transported, trimerized VSV-G remaining in the ER, as observed during the 10 °C experiment (Figure 7A). The fact that it does not implies that perhaps I14 preferentially recognizes densely concentrated VSV-G trimers such as those present on the virion surface used as immunogen (33). When trimers build up in the ER to high concentrations, such as at 10 °C, they are labeled by I14, however ongoing export at the permissive temperature must keep trimer concentrations in the ER below the threshold for I14 detection via immunofluorescence.

Since I14 can be used to visualize and quantify VSV-G at ERES that are otherwise difficult to detect due to excess unpackaged VSV-G, we used this technique to characterize the kinetics of VSV-G_ts045_ initial appearance at ERES after shift to the permissive temperature. Figure 7G displays the kinetic curve of the appearance of VSV-G spots in the I14 channel normalized to total VSV-G from the GFP channel. VSV-G appears at spots within seconds of temperature shift, and reaches a near-maximal steady-state value within ∼1 minute. To identify the I14-labeled early transport intermediates in detail, we compared the I14 spots to endogenous Sec31A. As shown in Figure 7H, the I14 spots at 60 seconds following temperature shift frequently co-localize with Sec31A, though not every I14 spot was positive for this marker. Finally, to quantify the relative density of trimers at ERES for cells with activated ALG-2, we performed 60-second incubations at the permissive temperature followed by fixation and immunostaining. As shown in Figure 7I, depletion of IP3R-3 or peflin to activate ALG-2 caused significantly greater intensity of spotty labeling in the trimer channel in cells with similar VSV-G expression indicated by the GFP channel. When this effect was quantified, we found that depletion of IP3R-3 caused an increase of 50% relative to control in cargo intensity at ERES, while depletion of peflin produced an 80% increase (Figure 7J). Figure S2A demonstrates that there were no significant differences in cargo expression levels between the same cells quantified in Figure 7J, and Figure S2B shows a standard 10-min transport assay on parallel coverslips of cells on the same day as the assay in 7J. Altogether this experiment indicates that during transport, ALG-2 activation drives significantly greater concentration of cargo at ERES, an effect that could explain the higher transport rate.

### ALG-2 activation increases COPII sorting stringency through interactions with Sec31A

To understand the relationship between ALG-2, Sec31A, and cargo sorting, we employed a double-knockdown approach using IP3R-3 and Sec31A siRNAs and added multiple cargoes to the experiment. As shown in Figures 8A and B, for VSV-G_ts045_-GFP, depletion of IP3R-3 in otherwise normal cells caused increased ER-to-Golgi transport, as previously documented. In Sec31A-depleted cells, however, IP3R-3 depletion caused no significant change in transport. To the contrary for the bulk-flow cargo GFP-FM_4_-hGH, depletion of IP3R-3 in otherwise normal cells caused a decrease in cargo trafficking, but in Sec31A-depleted cells, IP3R-3 depletion caused no significant change in transport. Thus, neither COPII client nor bulk-flow cargo are affected by IP3R-3 depletion when ER export is operating without Sec31A. Together with Figure 6A&B, this experiment demonstrates that the effects of basal Ca^2+^ signaling on transport require both Sec31A and ALG-2. In the same cells that were assayed for VSV-G or hGH transport, we collaterally assayed the intensity of Sec31A immunofluorescence. When Sec31A is recruited from the cytosol to ERES, the mean cytoplasmic Sec31A intensity increases due to greater concentration of the antigen at ERES. As shown in Figure 8C, Sec31A recruitment in cells expressing VSV-G increased significantly upon IP3R-3 depletion (bars 1 vs. 2). However, in cells expressing hGH, we detected no change in Sec31A (bars 5 v. 6). Importantly, this indicates that the presence of client cargo affects whether ALG-2 activation impacts Sec31A recruitment/retention at ERES. ALG-2 activation does not result in extra Sec31A recruitment unless an abundant client cargo is present, indicating that ALG-2 may enhance the mechanism by which the inner and outer coat sense and together retain cargo. We speculate that in the case of over-expression of the client or bulk-flow cargo constructs, the effects of endogenous cargos on Sec31A targeting were minimized.

**Figure 8.**
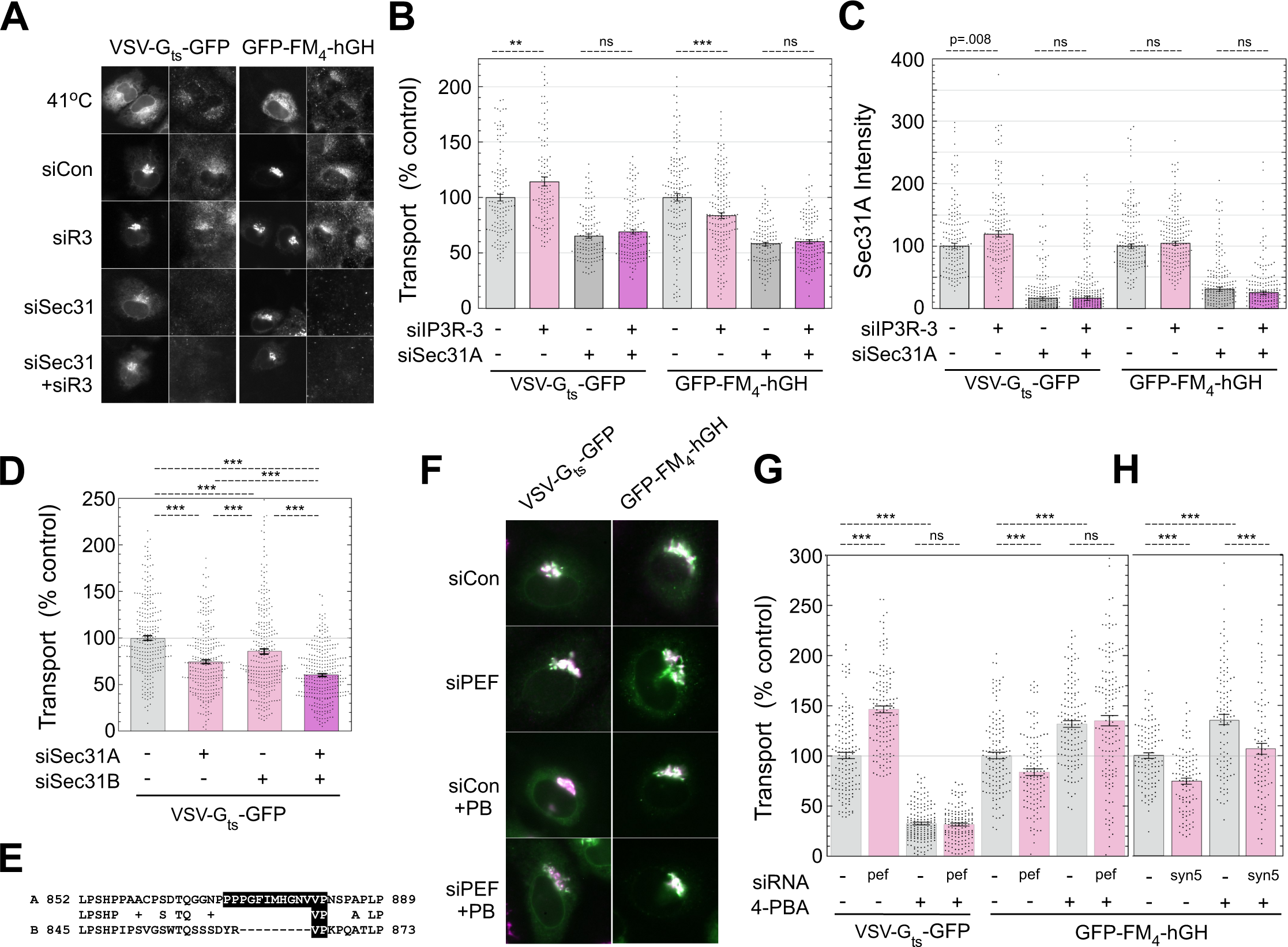
ALG-2 activation increases the stringency of COPII client cargo sorting through its interactions with Sec31A. **(A)** NRK cells were transfected with the indicated siRNAs and either VSV-G_ts_-GFP or GFP-FM_4_-hGH cargos, and subjected to a 10-min ER-to-Golgi transport assay, and immunolabeled for Sec31A. Shown are the GFP channel (left panel of each pair) and the Sec31A channel (right). **(B)** Quantitation of ER-to-Golgi transport as in Figure 5B. **(C)** Quantitation of whole-cell Sec31A intensity from the same cells quantified for transport in (B). **(D)** *Left,* quantified 10-min ER-to-Golgi transport assay on cells transfected with the indicated siRNAs and VSV-G_ts_-GFP cargo. **(E)** Alignment of rat Sec31A and Sec31B surrounding the necessary and sufficient ALG-2 binding site for Sec31A (black shaded residues). **(F)** NRK cells were transfected with control or peflin siRNAs as indicated and VSV-G_ts_-GFP or GFP-FM_4_-hGH cargos. 10-min transport assays were conducted in the presence or absence of 10 mM 4-PBA as indicated. Shown are merged images of the GFP and mannosidase II as in Figure 5A. **(G)** Quantitation of the transport assay shown in (F). **(H)** A follow-up control transport assay in which syntaxin 5 siRNA was substituted for peflin siRNA to illustrate specificity of 4-PBA for the sorting step in transport. p-values for two-tailed Student T-tests with unequal variance are indicated above plots; * = p ≤ .05; ** = p ≤ .005; *** = p ≤ .0005. Error bars show SEM.

It was somewhat unexpected that depletion of Sec31A, though very inhibitory to VSV-G transport, did not block it more completely. It has been found in yeast that expression of proteins that render the membrane more deformable can suppress the requirement for the COPII outer shell (35). We also showed in NRK cells that ≥95% depletion of Sec31A with siRNA results in a 40% increase in expression of sec23 (2), an essential member of the COPII inner coat. On the other hand, Sec31B could account for at least some of the residual transport. Indeed, we found that Sec31B was expressed in NRK cells and that its depletion caused a 15% inhibition of transport (Figure 8D). When combined with Sec31A depletion–which in this experiment caused 25% inhibition–Sec31B depletion caused an additional 15% inhibition, indicating that Sec31B contributes to transport in the absence of Sec31A. Immunoblots and additional siRNAs are shown in Figure S2 C-E. Knockdown of either Sec31 isoform caused over-expression of the other isoform, and KD of Sec31B strongly affected the mix of splicing isoforms of Sec31A. The lack of effects on transport by IP3R-3 depletion when Sec31A is also depleted (Figure 8B, bars 3 vs. 4 and 7 vs. 8) indicates that ALG-2 activation may not impact Sec31B. As shown in Figure 8E, Sec31B indeed lacks conservation to rat Sec31A residues 870-882, which are nearly identical to human residues 839-851, the necessary and sufficient ALG-2 binding site (7). Hence, a lack of effects of ALG-2 activation on Sec31B-supported transport is in fact predicted. Another conclusion from the experiment in Figure 8A and B is that both VSV-G and hGH rely upon Sec31A to the same degree for ER-to-Golgi transport, despite their undergoing opposite changes in transport upon IP3R-3 depletion (Figure 8B, bars 1 vs. 3 and 5 vs. 7). This demonstrates that the bulk flow cargo follows the same Sec31A-dependent pathway for transport that client cargo does. However, upon ALG-2 activation, COPII “chooses” VSV-G to the detriment of hGH. There is growing evidence that sorting involves the physical exclusion of non-client-cargo due to molecular crowding (36, 37). The higher Ca^2+^ environment does not merely enhance Sec31A function generally, leading to more vesicle transport of all cargoes, but rather increases the sorting stringency of COPII for client cargoes.

To more directly address the role of ALG-2 in COPII cargo sorting, we employed the drug 4-phenylbutyrate (4-PBA), which has been shown to specifically occupy the cargo-binding B-site on Sec24, occluding client cargo to cause client cargo-selective inhibition of budding (37). We reasoned that if ALG-2 activation were primarily affecting cargo sorting, then 4-PBA should neutralize any ALG-2 effects on ER-to-Golgi transport. We performed our standard transport assay and used peflin depletion to activate ALG-2. As shown in Figure 8F and G, peflin depletion produced the expected results on transport in the absence of 4-PBA; it stimulated VSV-G-GFP transport by 50% (part G, bars 1 vs. 2) and inhibited GFP-FM_4_-hGH transport by 20% (bars 5 vs. 6). Interestingly, the addition of 4-PBA to otherwise wildtype cells caused the opposite reciprocal results, with 4-PBA inhibiting VSV-G transport by 70% (bars 1 vs. 3) and stimulating hGH transport by 30% (bars 5 vs. 7). Stimulation of bulk-flow ER-to-Golgi transport by 4-PBA is predicted based upon its ability to cause secretion of ER resident proteins (37) and its ability to rescue plasma membrane localization of ER-retained quality control substrates (38). However, to our knowledge, it has not been directly observed before in kinetic measurements of ER-to-Golgi transport. A recent study with several cargos demonstrated that 4-PBA selectively inhibited secretion to the medium of client cargos without effects on bulk-flow cargos (39). Whether the stimulation of bulk flow is observed in any given experimental system may depend upon the particular cell type’s secretory capacity, intensity of competition between endogenous client and bulk flow cargoes, as well as the expression level and secretory efficiency of the model bulk flow cargo.

Finally, in Figure 8G bars 3 vs. 4 and 7 vs. 8, we found that peflin depletion had no significant effect on transport of either cargo in the presence of 4-PBA. Thus, when cargo sorting at the B site on Sec24 is blocked, ALG-2 activation cannot alter secretion of any cargo. This eliminates many other potential roles for ALG-2 in transport. For example, though ALG-2 works through the outer shell component Sec31A, it must not significantly affect the rate of vesicle budding, per se, since this would still affect transport in the presence of 4-PBA. To illustrate the specificity of this result, we subjected our bulk-flow cargo to inhibition by depletion of syntaxin 5, an ER/Golgi SNARE required for homotypic fusion of COPII vesicles (40). As shown in Figure 8H bars 1 vs. 2, syntaxin 5 depletion significantly inhibited ER-to-Golgi transport, as expected. Furthermore, as predicted, it also significantly inhibited transport in the presence of 4-PBA (bars 3 vs, 4), demonstrating that the experimental paradigm can distinguish distinct roles for ERES components. In conclusion, ALG-2 activation, caused by changes in basal Ca^2+^ oscillations or other perturbations, primarily acts to increase the stringency of client cargo sorting by the COPII coat.

## DISCUSSION

### Mechanism of effects on ER-to-Golgi transport

This work elucidates a role for background Ca^2+^ signals in setting the rate and specificity of ER export by modulating the activities and/or targeting of the PEF proteins ALG-2 and peflin at ERES. Specifically, this work discovered that ALG-2 activation drives higher concentrations of client cargo at ERES (Figure 7) and increases the stringency of COPII cargo sorting (Figure 8). These conclusions are based solely on functional transport measurements in intact cells, and will hopefully stimulate structural investigations to understand precisely how ALG-2 influences COPII cargo sorting. As noted earlier, the ALG-2 binding site on Sec31A (human Sec31A residues 839-851; (5)) is just N-terminal to the “active peptide” region (residues 981-1015) containing Trp-995 and Asn-996 that insert into the Sar1 active site to potentiate the Sec23 GAP activity that contributes to sorting (41, 42). We here note two distinct and non-exclusive mechanisms by which ALG-2 could regulate sorting. 1) ALG-2, in binding the PRR region, changes its conformation to render higher affinity interactions with the inner coat. Experimental support for this includes a previous study that found ALG-2 dramatically increased physical interactions between the outer and inner coat without directly bridging them (43). Since that study involved purified COPII components and ALG-2, there were no other ERES sites to which ALG-2 could have been anchored or targeted. 2) ALG-2 could help deliver, restrain or “place” the active peptide in the vicinity of Sar1, thus increasing its local concentration to favor more frequent interactions with the inner coat pre-budding complex causing more frequent Sar1 GTPase cycles. In this model ALG-2 acts as an adaptor between the PRR region and another ERES component. Experimental support for this includes frequent observations that ALG-2 stabilizes Sec31A on the membrane (4, 7, 8, 43) and the observations that ALG-2 can target an inhibitor peptide, comprised of an isolated ALG-2 binding region to an apparently saturable site at ERES to block ER export (2). A variant of the adaptor mechanism that apparently satisfies all of the experimental observations noted would be that ALG-2 homodimers crosslink the Sec31A PRR to other Sec31A PRR molecules, possibly nucleating a denser outer coat or alternatively making PRRs more potent or responsive to cargo by clustering them.

The precise mechanism of activation of ALG-2 by Ca^2+^ is still under investigation, though in vitro studies suggest that increased Ca^2+^ signals cause formation of ALG-2 homodimers at the expense of the ALG-2/peflin heterodimer (9) and favor homodimer binding to Sec31A over heterodimer; this explanation fits the prevailing concept that high Ca^2+^ favors binding of the ALG-2 homodimer to its various effectors (44). However, we also uncovered that decreases in peflin and ALG-2 protein abundance occur when IP3Rs are depleted (Figure 3). This suggests that Ca^2+^ may in addition regulate the species at ERES through altered stability or turnover of distinct PEF protein complexes. This hypothesis is supported by previous *in vitro* observations that heterodimer formation stabilizes both proteins against rapid proteasomal degradation (10), but that high Ca^2+^ causes dissociation of the heterodimer (9). A Ca^2+^-dependent turnover mechanism of removing inhibitory heterodimers would also be consistent with the relatively slow onset of secretion changes following agonist-induced Ca^2+^ signaling, first detectable after 30 minutes and increasing over several hours (8). ALG-2 and peflin have also been implicated in Ca^2+^-dependent ubiquitylation reactions at the ERES. Specifically, they were found to associate with Cul3^KLHL12^ to ubiquitylate Sec31A to produce enlarged COPII structures in cells (45, 46).

Both ALG-2 and peflin were required for the ubiquitylation which was proposed to be required for collagen secretion. However, neither ALG-2 nor peflin are required for secretory trafficking and peflin depletion *increases* collagen ER-to-Golgi transport in NRK cells (8). Furthermore, inhibition of CUL3 ubiquitylation activity with neddylation inhibitors did not affect collagen secretion, but instead CUL3^KLHL12^ appeared to regulate ER-lysosome trafficking for degradation (47). This also fits with recent observations that COPII and KLHL12 co-localized with procollagen at sites of non-canonical ER-phagy (48). Thus, CUL3^KLHL12^ ubiquitylation of Sec31A does not appear to be acutely involved in production of COPII vesicles. However, this does not rule out a role for ubiquitylation in regulation of cargo export from the ER, potentially involving ALG-2 and/or peflin as adaptors. Sec23 de-ubiquitylation has a rate-limiting impact on secretion in yeast though a ubiquitylation mechanism was not identified (49).

### IP3Rs and steady-state Ca^2+^ signals regulate protein secretion from the ER

Here we demonstrated that unexpectedly, IP3R-dependent Ca^2+^ signals are a rate-limiting factor setting the basal secretion rate in resting epithelial cells under normal growth conditions. This adds to the previously reported role of IP3Rs in secretion, in which agonist-dependent activation of IP3Rs can either increase or decrease the secretion rate depending upon the intensity and duration of signaling (8). The new data demonstrates that periodic, spontaneous Ca^2+^ oscillations, facilitated in part by intercellular Ca^2+^ waves (ICWs), provide a stimulus that determines how much ALG-2 and peflin are bound at ERES, which alters COPII targeting, sorting stringency, and ER export.

One consequence of this work is that physiological factors known to affect Ca^2+^oscillations or ICWs, such as Bcl-2 family protein expression (50), presenilin-1 mutation (51), or gap junctions, hemichannel function, and purinergic receptor expression (21), may also regulate the secretion rate through this mechanism. Furthermore, we demonstrated that this phenomenon is sensitive to expression levels and/or the mix of isoforms of IP3Rs present, with lower expression levels of IP3R-3 isoform favoring more spontaneous signaling and higher secretion rates. Other Ca^2+^ channels known to regulate IP3Rs could also regulate secretion by modulating IP3R-driven oscillations. Other factors that affect IP3R expression and turnover include micro-RNAs (52, 53), phosphatase and tensin homolog (PTEN) (54), BRCA1 associated protein (BAP1) (55), nuclear factor erythroid 2–related factor 2 (NRF2) (56), and NF-κB (57). These factors could potentially regulate the secretory export rate through the mechanism presented here.

Physiological phenomena known to be driven by spontaneous Ca^2+^ signaling could have changes in ER export as part of their mechanism. These phenomena could include glial control of neurovascular diameter (58, 59), nephron formation during kidney development (60), and numerous aspects of neural proliferation, migration, and differentiation during neocortical development (58, 61). While these phenomena involve spontaneous signaling, in most cases it is not known whether ICWs *per se* play a functional role. Retinal epithelial pigment cells are one cell type where ICWs have been established to regulate cell fate, influencing retinal development (21). Regulation of secretory output could be a significant effector mechanism by which ICWs alter cell fates in retinal or other cell types with ICWs. The biosynthetic secretory pathway is a rate-limiting determinant of plasma membrane and organelle biogenesis, plasma membrane composition, secretion of signaling molecules, and cell growth. Additionally, diseases associated with changes in IP3R expression, for example spinocerebellar ataxia (52), atherosclerosis (62), various liver disorders (63), and multiple cancer types including uveal melanoma, mesothelioma (64), colorectal carcinoma (65), and breast cancer (66, 67), could also have a secretory component to their pathophysiology. The Ca^2+^ channel polycystin-2 (also called TRPP2) interacts with and regulates the IP3Rs (68–70), and its dysfunction and disease pathology has been associated with excess constitutive secretion, in this case of extracellular matrix proteins (71). However, it should be noted that we have not tested whether other cell types besides NRK exhibit the strong link between spontaneous Ca^2+^ signaling and the secretion rate. Also note that “spontaneous” and “steady-state” are operational terms. We refer to spontaneous Ca^2+^ signaling as that which occurs in unperturbed cells growing in DMEM containing 10% FBS. The initiation of the signals could involve trace agonists, for example serotonin present in typical lots of FBS (72).

### Roles of IP3Rs in regulating the luminal Ca^2+^ store

The fact that IP3R depletion caused a mild reduction of luminal Ca^2+^ is challenging to interpret, since store-operated Ca^2+^ entry (SOCE) should be more than fast enough to compensate for the periodic Ca^2+^ oscillations we observed in resting NRK cells. One interpretation would be that IP3Rs play a specific role in SOCE in NRK cells. Indeed, a direct role of IP3Rs in SOCE in *Drosophila* has been demonstrated (73); however, their direct role in SOCE in mammalian cells, though widely discussed, is not well-supported (74). Though we did not pursue the mechanism of the effects on Ca^2+^ stores, control experiments in Figure 6 allowed us to determine that the changes in luminal Ca^2+^ caused by IP3R siRNAs did not drive the effects we observed on ER-to-Golgi transport.

## EXPERIMENTAL PROCEDURES

### Antibodies

Mouse monoclonal anti-mannosidase II antibody was purchased from Covance Research Products (Denver, PA; product MMS-110R-200). Rabbit polyclonal anti-IP3R-1 antibody was purchased from ThermoFisher Scientific (Waltham, MA; product PA1-901). Mouse monoclonal anti-IP3R-3 antibody was purchased from BD Biosciences (San Jose, CA; item 610313). Anti-rbet1 mouse monoclonal antibody, clone 16G6, was as described (26). Rabbit polyclonal anti-peflin and chicken polyclonal anti-ALG-2 were produced previously (8). Rabbit polyclonal anti-sec31A, anti-sec23, anti-p24 and anti-β-Cop were produced previously against synthetic peptides (1, 40, 75). Rabbit polyclonal anti-Phospho-eIF2α antibody was purchased from Cell Signaling Technology (Danvers, MA; product 9721s). Rabbit polyclonal anti Phospho-IRE1 was purchased from Abcam (Cambridge, United Kingdom; item ab48187). Rabbit polyclonal anti-ATF4 antibody was purchased from GeneTex (Irvine, Ca^2+^; product GTX101943). Rabbit polyclonal anti-sec24c was a kind gift from Dr. William Balch (Scripps Institute). Rabbit polyclonal anti-p115 was produced in-house (76). Rabbit polyclonal anti-p58 was a kind gift from Dr. Jaakko Saraste (University of Bergen, Norway). I14 conformation-specific anti-VSV-G_ts045_ monoclonal antibody was kindly provided by Dr. John Ngsee (University of Ottawa). Secondary antibody conjugates with Alexa Fluor^TM^ 488 were from Invitrogen (Carlsbad, CA; product A11001). Cy3- and cy5-conjugated secondary antibodies were purchased from Jackson ImmunoResearch Laboratories (West Grove, PA).

### qRT-PCR

NRK cells were transfected with IP3R-3 siRNA (see below) using Transfast (Promega Corp.; Madison WI) using manufacturer’s instructions. Total RNA was isolated using the PEQLAB total RNA isolation kit (Peqlab; Erlangen, Germany) and reverse transcription was performed in a thermal cycler (Peqlab) using a cDNA synthesis kit (Applied Biosystems; Foster City, CA). mRNA levels were examined by qRT-PCR. A QuantiFast SYBR Green RT-PCR kit (Qiagen; Hilden, Germany) was used to perform real time PCR on a LightCycler 480 (Roche Diagnostics; Vienna, Austria), and data were analyzed by the REST Software (Qiagen). Relative expression of specific genes was normalized to human GAPDH as a housekeeping gene. Primers for real time PCR were obtained from Invitrogen (Vienna, Austria).

### siRNA knockdowns and transfections

NRK cells were electroporated with 0.6 µM siRNA and grown in DMEM with 4.5 g/L glucose containing 10% fetal calf serum and 1% penicillin-streptomycin. After 2-3 days of normal growth at 37 °C, the cells were resuspended and re-electroporated, this time with a combination of the siRNA plus 7.5 µg of protein plasmid. Cells were allowed to recover and grow on coverslips at 37 or 41 °C for 24 h.

Alternatively, sub-confluent cells grown on glass coverslips in 6-well plates were directly transfected without resuspension. After ∼16 h incubation in OptiMEM containing 0.3% RNAiMax (Invitrogen; Carlsbad, CA) and 0.28 µM siRNA, transfection medium was supplemented with an equal part of OptiMEM containing 5% fetal calf serum, 2 mM CaCl_2_, and 1% penicillin-streptomycin and incubated for another 3-5 hours. Subsequently, this post-transfection medium was removed (but saved) and replaced with DMEM containing 10% fetal calf serum, 1% penicillin-streptomycin, 0.3% PolyJet (SignaGen Laboratories; Frederick, MD), and 1 µg/ml plasmid. After ∼12 h incubation, the plasmid solution was removed, cells were washed twice with PBS and returned to their siRNA-containing OptiMEM medium at 37 or 41 °C to recover while continuing the knockdown for another 24 h. There was no noticeable difference in knockdown or transfection efficiency between the two transfection protocols. Cells from all transfections were either lysed directly in SDS-PAGE sample buffer for quantitative immunoblotting, or else processed for transport, colocalization, or FRET assays as described below.

Plasmid constructs GFP-VSV-G_ts045_, GFP-FM_4_-VSVG_tm_, and GFP-FM_4_-hGH were described previously (8). RUSH-VSV-G-GFP was purchased from Addgene (Watertown, MA; catalog # 65300). D1ER (22) and D3cpv (23) were described previously.

Custom siRNAs were purchased from Ambion/Thermo Fisher (Waltham, MA) or Gene Link (Elmsford, NY) and contained no special modifications. We used two different siRNAs for IP3R-3 which appeared to perform similarly in knockdown efficiency and effects on Ca^2+^ and secretion; the large majority of experiments were performed using siRNA Itpr3-7848 (5’ GCA GAC UAA GCA GGA CAA A dTdT 3’). siRNA Itpr3-0452 (5’ GGA UGU GGA GAA CUA CAA A dTdT 3’) was used intermittently for confirmation purposes. ALG-2 and Peflin siRNAs were described and their effects on secretion confirmed previously (2, 11). Sec31A siRNA had the following sense strand sequence: 5’ GAC CUU UGU UUA CAC GAU dTdT 3’. Sec31B siRNA had the following sense strand sequence: 5’ CCU UGA UUG CCC AGA AAC A dTdT 3’ or 5’ GGA AAU CUC CUU AAU UAU U dTdT 3’. Syntaxin-5 siRNA had the following sense strand sequence: 5’ CCA UUG UAG UUC AUU GCA dTdT 3’. Control siRNAs were custom synthesized non-targeting siRNAs from the same manufacturer. Immunoblotting of cell lysates enabled validation of knockdown efficiencies for each siRNA experiment that was functionally analyzed.

### ER-to-Golgi transport assay

NRK cells were transfected with cargo protein plasmid and siRNA as described above and plated on poly-L-lysine coated coverslips. For VSV-G_ts045_, cells were shifted to 41 °C for 6 to 15 h prior to transport, to accumulate the cargo in the ER. For the transport assay, cells were either fixed by dropping coverslips directly into fixative or into 6-well transport chambers with pre-equilibrated 32 °C medium for 10 min prior to transfer to fixative. Alternatively, for FM4-containing cargo constructs and RUSH-VSV-G, transfected cells were kept at 37 °C, and coverslips were fixed either by dropping coverslips directly into fixative or into 6-well transport chambers for 10 min prior to transfer to fixative. In this case, transport chambers contained 37 °C media containing either 500 nM AP21998, also known as D/D solubilizer (TakaraBio: 635054), for FM4-containing cargos; or 40 µM Biotin for RUSH cargos. Coverslips were fixed and labeled using anti-mannosidase II antibody as described in the immunofluorescence section below. For transport experiments using 4-phenylbutyrate (4-PBA) as shown in Figure 8 G-H, transfected cells were incubated in media with 10 mM 4-PBA for 1 hour prior to transfer to transport chambers also containing 10 mM 4-PBA. 4-PBA was kindly provided by Dr. Michael Kavanaugh (University of Montana). For live-cell transport assays as shown in Figure 5 C-E, coverslips were placed in a perfusion chamber and imaged on a Nikon TE300 inverted microscope equipped with a 40x objective, Crest Optics X-Light v1 spinning disk scanner, 89 North LDI-7 laser launch, and a PCO Edge 4.2 sCMOS camera, all automated with Micro-Manager software. Imaging was carried out using a constant perfusion rate of 2 ml/min maintained at 37 °C with imaging at 60-second intervals. After 2 min of DMEM, the perfusion was switched to DMEM + 500 nM D/D solubilizer and imaged for another 20 min.

Morphological quantitation of ER-to-Golgi transport was accomplished as described previously (8, 11). Images were collected in a consistent manner with regard to cell morphology, protein expression levels and exposure. A single image plane was collected for each color channel (GFP and mannosidase II) for each field of cells randomly encountered; image deconvolution was not performed. In most cases widefield images were used (microscope is described below in immunofluorescence section), while in others (Figure 5C-E and Figure 7A-J), spinning disk confocal images were used for quantitation of transport with similar results.

Images were analyzed using ImageJ open source image software with automation by a custom script (available upon request) that randomizes the images and presents them to the user in a blind fashion. The user identifies transfected cells and subcellular regions while the script extracts parameters including extracellular background, Golgi intensity and diffuse cytoplasmic ER intensity. Transport index is calculated for each individual cell as (Golgi_max_ - background) / (ER_mean_ - background), and all extracted parameters are written to an appendable output file along with a cell number and image title so that the data was traceable. Using this method, the user quantitates about 60 cells per hour. Once transport indices have been obtained for all conditions in an experiment, each value is subtracted by the mean transport index value for cells that were transferred directly to fixative without a transport incubation (typically a value between 1.0 and 1.5) to generate the net transport index. Net transport indices are then normalized to the mean siControl value for the particular experiment, prior to plotting and comparison between experiments. Each result reported here was obtained in at least three separate experiments on different days.

For 10 °C ERES loading experiments as shown in Figure 7 A-B, cells were transfected with VSV-G_ts045_ and prepared as described above, then either fixed by dropping coverslips directly into fixative or into 6-well transport chambers with pre-equilibrated 10 °C medium for 1 hour prior to transfer to fixative. Quantitation of GFP spot prominence was accomplished using a custom ImageJ script (available upon request). Briefly, GFP spots are identified visually and a compound ROI is created around each spot. The compound ROI consists of a circular inner ROI fully encompassing the spot and a larger circular outer ROI encompassing the inner ROI. The script then extracts parameters including extracellular background, inner ROI intensity, and outer ROI intensity. The outer ROI intensity measurement excludes the area comprising the inner ROI, thus specifically measuring GFP intensity directly adjacent to the identified spot. Spot prominence is calculated for each individual cell as (Inner ROI_max_ – background) / (Outer ROI_mean_ – background).

### Immunofluorescence microscopy

Coverslips were fixed with 4% paraformaldehyde containing 0.1 M sodium phosphate (pH 7.0) for 30 min at room temperature and quenched twice for 10 min with PBS containing 0.1 M glycine. Fixed cells were treated for 15 min at room temperature with permeabilization solution containing 0.4% saponin, 1% BSA, and 2% normal goat serum dissolved in PBS. The cells were then incubated with primary antibodies diluted in permeabilization solution for 1 h at room temperature. Next, coverslips were washed 3x with permeabilization solution and incubated 30 min at room temperature with different combinations of Alexa Fluor^TM^ 488-, Cy3-, and/or Cy5-conjugated anti-mouse, anti-rabbit, or anti-chicken secondary antibodies. After the secondary antibody incubation, coverslips were again washed 3x using permeabilization solution and mounted on glass slides using Slow Fade Gold antifade reagent (Invitrogen product S36936) and the edges sealed with nail polish. Slides were analyzed using a 40x or 60x objective on a Nikon E800 widefield microscope with an LED illumination unit (CoolLED pE 300^white^), PCO Panda sCMOS camera, Prior excitation and emission filter wheels and Z-drive, all automated using Micro-Manager software.

### Colocalization Assays

For immunofluorescence co-localization experiments, fixed NRK cells such as in Figure 4 were captured using a 60x/1.40 Plan Apo objective as z-stacks in eleven 200-nm increments for each color channel. Image stacks were deconvolved using Huygens Essential Widefield software (Scientific Volume Imaging, Hilversum, The Netherlands). Final images for display and quantitation represent maximum intensity projections of deconvolved stacks. ERES area was assessed by a custom ImageJ script (available upon request). Briefly, background labeling was removed by defining a dark extracellular area of each channel image as zero. For each cell an object binary image mask (excluding the nucleus) was generated for each individual channel by thresholding using the ‘Moments’ (for rbet1) or ‘Renyi Entropy’ (for VSVG, ALG-2, Peflin, and sec31a) algorithms. Next, each channel mask was compared to the other using a Boolean ‘and’ operation to create co-localization binary masks. Individual channel masks and co-localization masks were converted into ROIs and applied to each original channel image to measure total area and fluorescence intensity within the area specified by the mask. All measurements were extracted and written to an appendable output file along with the cell number and image title so that the data was traceable. Colocalization area and fluorescence intensity measurements for each cell were then normalized to the mean siControl value for the particular experiment, prior to plotting and comparison.

### Calcium Imaging

Subconfluent NRK cells growing on glass coverslips were transfected with genetically encoded Ca^2+^ sensors D1ER or D3cpv and siRNA as described above and imaged the next day. In some experiments (Figures 1C-F and 2) control or knockdown cells were loaded with FURA-2-AM for 30 minutes at 37°C directly prior to imaging, rather than transfection with a Ca^2+^ sensor. Coverslips were placed in a perfusion chamber and imaged on a Nikon TE300 inverted microscope equipped with a 40x objective, motorized high speed Sutter Lambda filter wheel for emissions, CoolLED pe340 excitation system, and PCO Panda sCMOS camera, automated with Micro-Manager software. Imaging was carried out using various perfusion protocols as described below with imaging at 3 or 4-second intervals. For examination of basal Ca^2+^ oscillations as in Figure 2, cells were labeled with FURA-2AM and imaged for 20 minutes in non-perfusing DMEM containing 10% FBS maintained at ∼37°C. For examination of agonist-dependent Ca^2+^ signaling as in Figure 1C-F, cells were labeled with FURA-2AM and imaged for a total of 10 minutes with a constant perfusion rate of 2 ml/min at room temperature. After 5 minutes of DMEM, the perfusion solution was changed to DMEM + 1 µM bradykinin. Each FURA imaging interval involved collecting an image at 510 nm emission and 340 or 380 nm excitation. For CFP/YFP FRET Ca^2+^ sensors, each imaging interval involved collecting images at 480 nm and 530 nm using 430 nm excitation. For examination of ER luminal Ca^2+^ stores as in Figures 6C-E and S1A, cells were transfected with D1ER and imaged for a total of 8 minutes with a constant perfusion rate of 2 ml/min at room temperature. After 5 min of 2Ca^2+^ buffer (2 mM CaCl2, 138 mM NaCl, 1mM MgCl2 5 mM KCl, 10 mM D-glucose, 10 mM Hepes, pH 7.4), the buffer was changed to 0Ca^2+^ buffer (138 mM NaCl, 1 mM MgCl2, 5mM KCl, 10 mM D-glucose, 0.1 mM EGTA, 10 mM Hepes, pH 7.4) + 3 µM Ionomycin. For examination of cytosolic Ca^2+^ release as in Figure S1C, cells were transfected with D3cpv and imaged for a total of 15 minutes with a constant perfusion rate of 2 ml/min at room temperature. After 5 minutes, the 2Ca^2+^ buffer was changed to 2Ca^2+^ buffer + 200 nM Thapsigargin (Tg). For analysis in ImageJ, each cell, as well as an extracellular background region was enclosed in an ROI and mean intensity was collected in each color channel at each time interval. Data was imported to Kaleidagraph software, where the intensity data was converted to ratios. For FURA, R= (emission at 510 nm when excited at 340 nm - background at 510/340) / (emission at 510 nm when excited at 380 nm - background at 510/380). For FRET sensors, with excitation at 430 nm, R= (emission at 530 nm - background at 530) / (emission at 480 nm - background at 480). The FRET or FURA ratio curves were then converted to R/R0 by dividing every R value by the average of images 2-8 for each trace and, when necessary, fit to a polynomial or exponential decay function to remove effects of progressive photo-bleaching or baseline drift.

## ACKNOWLEDGMENTS

This work was supported by NIH grant 2R15GM106323-03 to J. C. H., and by the University of Montana Stella Duncan Memorial Fellowship to A.J.H. The authors thank Mr. Tyler Brown for expert assistance in the quantitation of Figure 5C-E.

---

**Figure S1.**
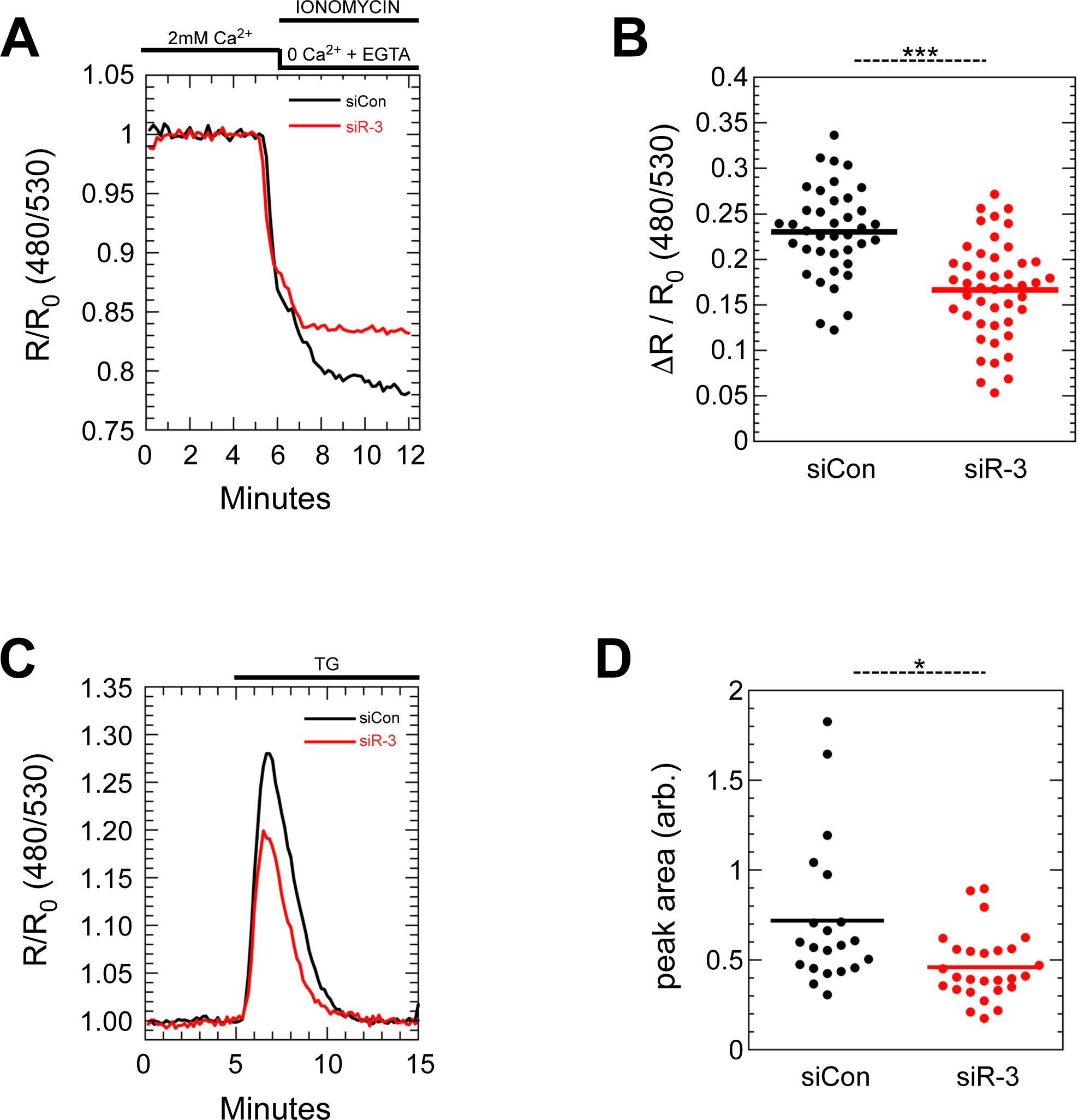
IP3R-3 depletion causes reduced ER luminal Ca^2+^ stores. **(A)** Representative traces of ER luminal Ca^2+^, using the sensor D1ER, upon depletion with ionomycin and low Ca^2+^ medium. **(B)** Quantitation of the luminal Ca^2+^ drops in cells from three experiments of Ca^2+^ depletion traces as in (A). **(C)** Representative traces of cytosolic Ca^2+^, using the sensor D3cpv, upon release of luminal Ca^2+^ stores by addition of thapsigargin to the extracellular medium. **(D)** Quantitation of cells from two experiments of the peak areas in traces as in (C). In (B) and (D), each dot represents one cell.

**Figure S2.**
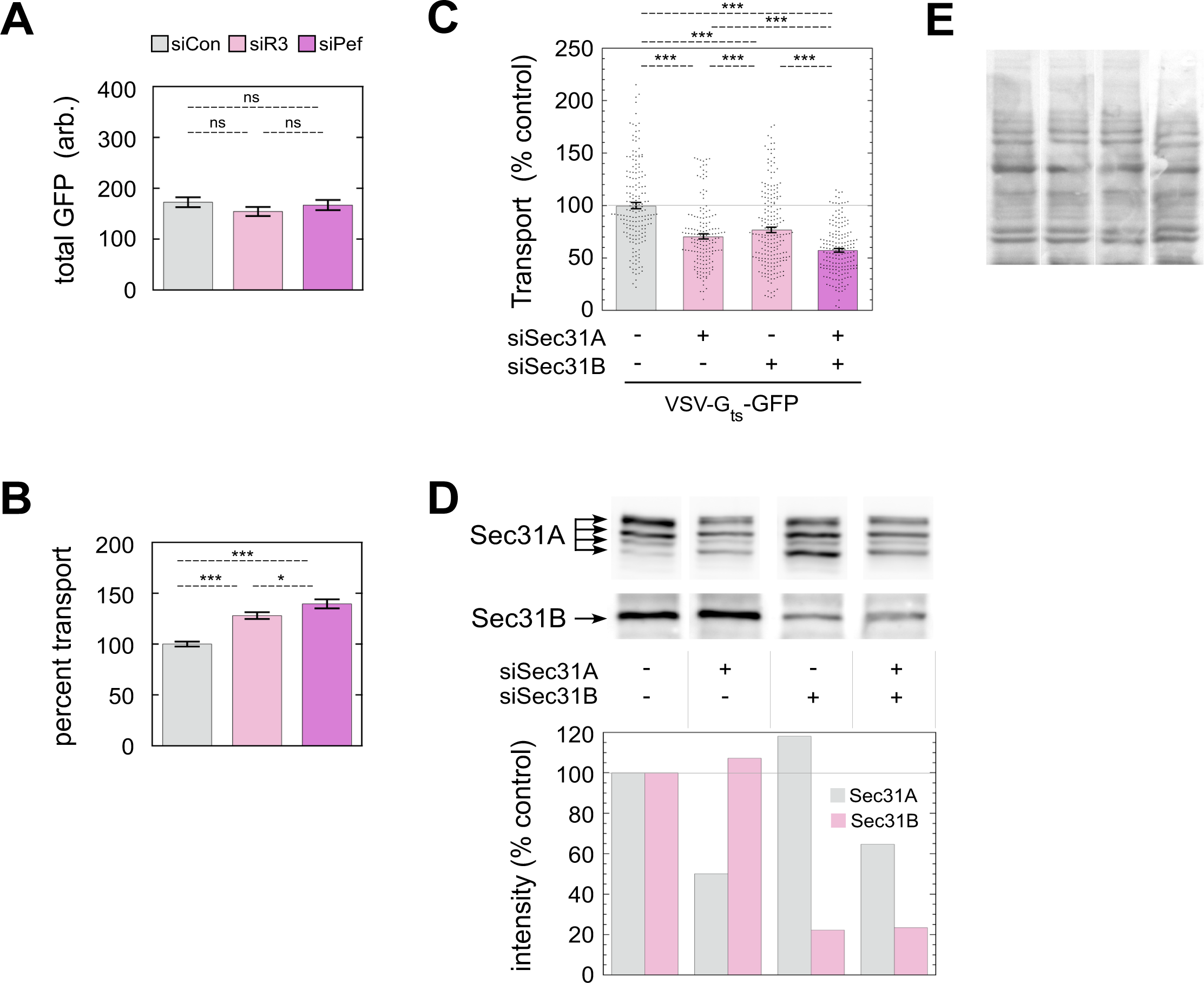
Additional controls for experiment in Figures 7J and 8D. **(A)** Quantitation of GFP intensity of the entire cytoplasmic area (nucleus excluded) of the same cells quantified in Figure 7J. **(B)** Standard 10-min transport assay on parallel coverslips from the same experiment as Figure 7J. **(C)** Repeat of transport assay from Figure 8D but using an entirely distinct Sec31B siRNA sequence. **(D)** Immunoblotting of cells from the experiment in Figure 8D (top) and quantitation of the blots (bottom). For Sec31A quantitation, all four indicated isoforms were included. The RefSeq database lists 17 predicted rat splicing isoforms of Sec31A, only the largest of which has been empirically documented. **(E)** Ponceau staining of the blot from (D) prior to immunoblotting. p-values for two-tailed Student T-tests with unequal variance are indicated above plots; * = p ≤ .05; ** = p ≤ .005; *** = p ≤ .0005. Error bars show SEM.

